# Cross-linkers at growing microtubule ends generate forces that drive actin transport

**DOI:** 10.1101/2021.07.09.451744

**Authors:** Celine Alkemade, Harmen Wierenga, Vladimir A. Volkov, Magdalena Preciado López, Anna Akhmanova, Pieter Rein ten Wolde, Marileen Dogterom, Gijsje H. Koenderink

## Abstract

The actin and microtubule cytoskeletons form active networks in the cell that can contract and remodel, resulting in vital cellular processes as cell division and motility. Motor proteins play an important role in generating the forces required for these processes, but more recently the concept of passive cross-linkers being able to generate forces has emerged. So far, these passive cross-linkers have been studied in the context of separate actin and microtubule systems. Here, we show that crosslinkers also allow actin and microtubules to exert forces on each other. More specifically, we study single actin filaments that are cross-linked to growing microtubule ends, using in vitro reconstitution, computer simulations, and a minimal theoretical model. We show that microtubules can transport actin filaments over large (micrometer-range) distances, and find that this transport results from two antagonistic forces arising from the binding of cross-linkers to the overlap between the actin and microtubule filaments. The cross-linkers attempt to maximize the overlap between the actin and the tip of the growing microtubules, creating an affinity-driven forward condensation force, and simultaneously create a competing friction force along the microtubule lattice. We predict and verify experimentally how the average transport time depends on the actin filament length and the microtubule growth velocity, confirming the competition between a forward condensation force and a backward friction force. In addition, we theoretically predict and experimentally verify that the condensation force is of the order of 0.1 pN. Thus, our results reveal a new mechanism for local actin remodeling by growing microtubules.

**Significance Statement:** Complex cellular processes such as cell migration require coordinated remodeling of both the actin and the microtubule cytoskeleton. The two networks for instance exert forces on each other via active motor proteins. Here we show that, surprisingly, coupling via passive cross-linkers can also result in force generation. We specifically study the transport of actin filaments by growing microtubule ends. We show by cell-free reconstitution experiments, computer simulations, and theoretical modeling that this transport is driven by the affinity of the cross-linker for the chemically distinct microtubule tip region. Our work predicts that growing microtubules could potentially rapidly relocate newly nucleated actin filaments to the leading edge of the cell and thus boost migration.

## Introduction

Vital cellular processes such as cell division and motility are driven by contraction and remodelling of active networks within the cell: the actin and microtubule cytoskeletons. These contraction and remodelings processes require the generation of forces and relative movement of filaments. It is well known that motor proteins can cross-link filaments, and organize the network by driving the relative movements of the filaments [1]. Besides the well-appreciated role of motor proteins, also polymerization dynamics have been shown to generate forces required for self-organization and remodeling [2–4]. In addition, the importance of passive (non-motor) cross-linkers has recently been increasingly recognized. It is now clear that passive cross-linkers can generate driving forces too, as has been shown for anillin in the contractile ring [5] and Ase1 in the spindle [6]. These forces are generated via the entropy associated with the cross-linker’s diffusion within the overlap region [6], or via the condensation (or preferential binding) of cross-linkers to the overlap between filaments [5, 6], therefore named ‘condensation force’. In addition, passive cross-linkers create frictional forces [6,7], which not only affect the speed at which cytoskeletal structures are remodeled, but can even be essential for their stability by opposing the motor proteins [8, 9].

The studies on force generation by passive cross-linkers have so far focused on individual cytoskeletal systems. However, passive cross-linkers could also be a way for two systems to exert forces on each other. In particular, there is a growing body of work showing the importance of microtubule/actin crosstalk in vital cellular functions [10]. Besides crosstalk via signalling pathways, it has been shown that crosstalk via motor proteins such as myosin-X (Myo10) is among the drivers of remodeling the microtubule spindle network during cell division [11,12]. In addition, various passive cross-linking proteins have been identified to enable crosstalk, such as the family of spectraplakins, which are proteins that can bind both actin and microtubules. The most prominent and best-studied member is ACF7 (MACF1) and its Drosophila homologue Short stop (Shot), which contain both microtubule lattice- and end-binding activities. These cross-linkers play a crucial role in a number of cellular processes, such as cell migration, cell-cell connections, vesicular transport, cell polarity and cell division [10, 13]. In addition to ACF7, also other passive cross-linking proteins such as tau and Gas2Like1 have been shown to result in co-alignment and stabilization of the actin and microtubule networks [14–17]. Another example of a microtubule end-binding cross-linker that links actin to microtubule ends via EB3-mediated interactions is drebrin, which is required for sprouting neurites from a neuronal cell [18].

Recent experiments suggest that the driving condensation forces and friction forces associated with passive crosslinkers can also couple to the growth dynamics of filaments, and thereby allow passive crosslinkers to play a central role not only in contraction but also in transport. It was shown that an engineered passive cross-linker called TipAct can mediate transport of actin filaments by binding to the tips of growing microtubules [15]. The microtubule tip region is chemically different from the microtubule lattice region, due to delayed hydrolysis of GTP that is associated with tubulin subunits that incorporate into growing microtubules. This region is recognized by microtubule end-binding (EB) proteins [19–21] as well as EB-binding partners such as TipAct. However, the processivity of the actin transport mechanism and its dependency on relevant parameters such as actin filament length and microtubule growth velocity, as well as the magnitude of the generated forces are currently unresolved.

Here, we investigate the conditions necessary for microtubule-mediated actin transport and elucidate the cross-linker-dependent mechanism. We combine biochemical reconstitution experiments with coarse-grained computer simulations and a minimal theoretical model, and show that in the presence of passive cross-linkers, microtubules can transport actin filaments over large (micrometer-range) distances during periods that last up to several minutes. We propose and test a new mechanism to explain this movement, in which actin transport is the result of a competition between a forward force that tends to maximize the overlap of the actin filament with the plus-end of a growing microtubule and a backward force caused by the friction between the actin and microtubule lattice. These two antagonistic forces are both caused by the binding of cross-linkers. The preferential binding of cross-linkers at the interface between two objects (such as filaments) creates a driving force that tends to maximize the overlap region. This type of force, which, following earlier work [6, 7], we will call condensation force, can drive the movement of intracellular cargoes [2, 22–24] and the contraction of filament bundles [5, 6, 25, 26] in various biological contexts. However, the combination of a processive transport of cargo by a cross-linker based condensation force towards a chemically distinct region that is simultaneously hindered by a friction force caused by the same cross-linkers is specific to filament transport and has to our knowledge not been studied before. The active transport of filaments by the growing microtubule plus-ends thus provides a new minimal mechanism by which two cytoskeletal systems can interact, which is distinct from cytoskeletal crosstalk that is driven by motor proteins [11, 27–30]. We expect this mechanism to be especially relevant in migrating cells, where growing microtubules nucleate actin filaments and encounter membrane-bound actin arrays.

## Results

### Growing Microtubules Plus-Ends Can Transport Actin Filaments

To study actin filament transport by growing microtubule plus-ends, we use the engineered crosslinking protein TipAct that contains both a domain that specifically binds actin filaments and a domain that recognizes end-binding (EB) proteins to target it to the growing ends of microtubules (Fig. S1A,B) [15]. Dynamic microtubules are grown from surface-immobilized seeds stabilized with the nucleotide analogue GMPCPP and incubated with fluorescently labeled tubulin. The microtubules grow in the presence of TipAct, EB3, and freely diffusible stabilized actin filaments and are imaged by total internal reflection fluorescence (TIRF) microscopy (Fig. 1A). We observe a 54μm×54μm sized region of interest, which typically contains on the order of 30 microtubules (Fig. 1B), and we look for instances where growing microtubules interact with actin filaments (see Materials&Methods for criteria). Figure 1C shows a typical example of an encounter of a growing microtubule with an actin filament. Once the microtubule starts growing, an EB3-TipAct complex localizes at its growing end (yellow arrow). After several seconds, an actin filament binds and co-localizes to this tip-tracking complex (blue arrows) and is transported over a distance of several micrometers, before finally unbinding while the microtubule continues growing (white arrows).

**Figure 1:**
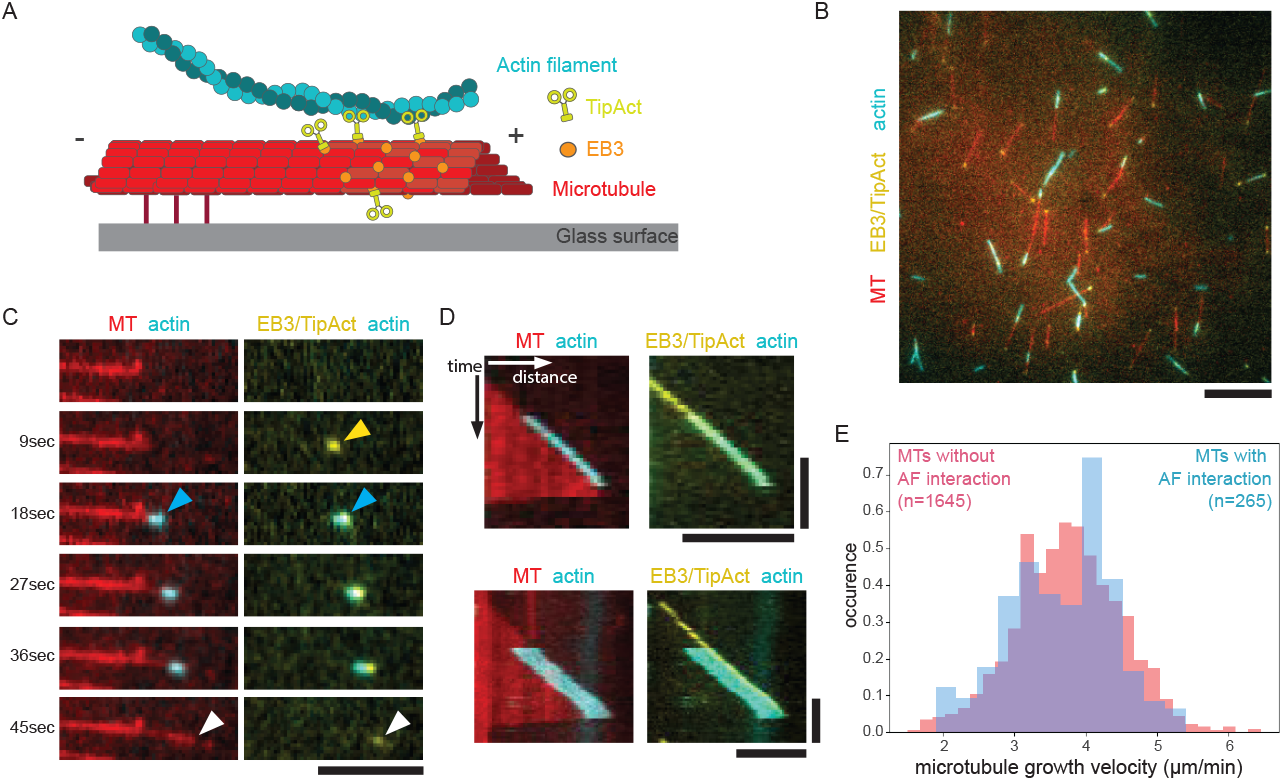
Actin transport by growing microtubule plus-ends. (A) Schematic of the experimental assay for observing microtubule-mediated actin transport, showing a stabilized actin filament (cyan), the engineered cytolinker TipAct (green-yellow), and the microtubule end-binding protein EB3 (orange) moving freely in solution, while the growing microtubule (red) is anchored to the surface of a functionalized and passivated glass slide. (B) Experimental field of view, showing a typical cropped region that was used for analysis, including microtubules/tubulin (red), EB3 (yellow), TipAct (yellow), and stabilized actin filaments (cyan). (C) Time series of the growing plus-end of a microtubule that recruits and transports an actin filament via EB3/TipAct complexes. Arrowheads show the localization of the EB3/TipAct-complex (yellow), the binding of a short actin filament (cyan), and the unbinding of this actin filament while the microtubule continues to grow (white). (D) Kymographs (space-time plots) of actin filament transport by the growing plus-end of a microtubule, showing the microtubule (red), EB3 and TipAct (yellow), and an actin filament (cyan). From these kymographs, we measure parameters of the microtubule dynamics, such as the growth velocity. (E) Normalized distribution of microtubule growth velocities for growth events where the microtubules does not interact with actin filaments (red) and for microtubules that transport an actin filament (blue). The average growth velocities are 3.5±0.6 μm/min and 3.7±0.7 μm/min for non-interacting and interacting events, respectively. Separate channels are shown in Fig. S1C-F. Both EB3 and TipAct are GFP-labelled. Scale bars: 10 μm in B, 5 μm in C, 5 μm (horizontal) and 60 s (vertical) in D.

To analyze the effect of actin transport on microtubule growth dynamics, we construct a kymograph, i.e. a space-time plot along the length of the microtubule (Fig. 1D). From the microtubule tip position as a function of time, we can extract the microtubule growth velocity. In the example kymographs in Fig. 1D, actin transport does not affect the growth velocity. This finding is confirmed by an analysis of 265 transport events (Fig. 1E).

To further analyze transport parameters such as the transport rate and duration, we again turn to kymographs. Fig. 2A shows an example kymograph corresponding to the transport event shown in Fig. 1C. It clearly visualizes the recruitment of the actin filament shortly after the microtubule starts growing, and its unbinding before the microtubule undergoes a catastrophe. For each event, we measure the transport time from the moment when the actin filament begins to move along with the growing microtubule tip region until this directed motion stops. Since EB3/TipAct can also tracking the growing minus-end of a microtubule (as visible in Fig. S3A), actin transport also occurs at these minus-ends. However, due to the greater physiological relevance, we focus on the microtubule plus-ends.

**Figure 2:**
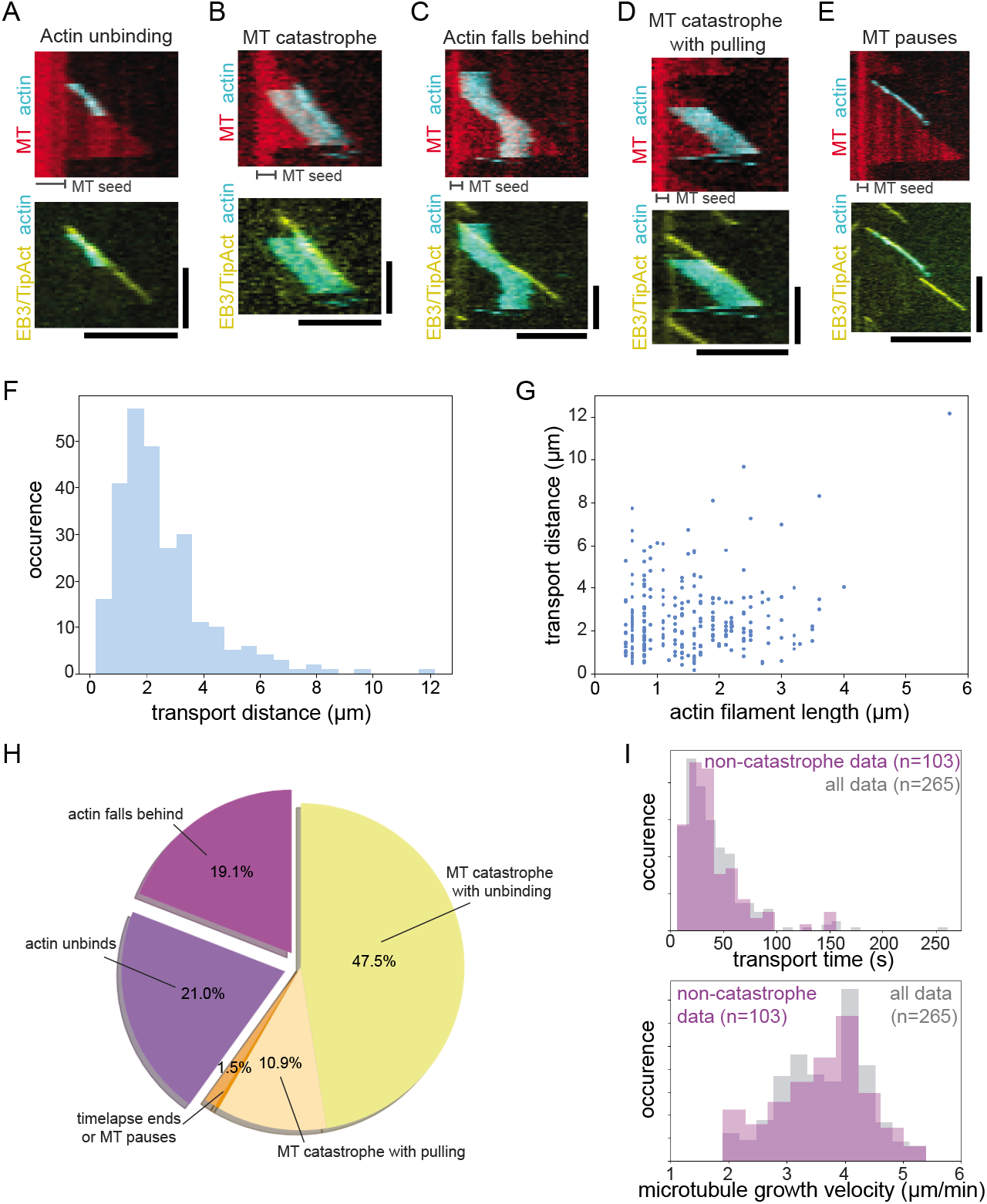
Mechanisms limiting the actin transport time. (A) Kymograph of the actin transport event depicted in Fig. 1C, showing that the event ends by the unbinding of the actin filament while the microtubule continues growing for a while (see also Movie S1). (B) Kymograph of a typical transport event that ends upon a microtubule catastrophe (see also Movie S2). (C) Kymograph of a typical transport event that ends by loss of contact of the actin filament with the tip resulting in the actin filament falling behind and lingering on the MT lattice (see also Movie S3). (D) Kymograph of a typical transport event that ends upon a microtubule catastrophe, but where the shrinking microtubule pulls an actin filament backwards (see also Movie S4). (E) Kymograph of the transport event that ends by the disappearance of the comet at a pausing microtubule. This is the only example of transport ended by microtubule pausing. (F) Distribution of transport distances for actin filaments transported by growing microtubule plus end. Median distance of 2.1 μm (and mean of 2.5 μm) and range of 0.8-12.2 μm. (G) Transport distance as a function of actin filament length. Actin filaments that were transported by the growing microtubule plus-end have a median length of 1.4 μm (and mean of 1.6 μm) and a length range of 0.5-7.9 μm. The maximal length is limited by the tendency of longer filaments to form bundles, which are excluded from further analysis. (H) Categories of termination events together with their observed frequency (n=265). Typical examples of the categories are shown in A-E. (I) Distributions for the transport times (top) and microtubule growth velocities (bottom) are indistinguishable when we consider the complete dataset (n=265) or the subset of transport events that end by the actin filament unbinding from the microtubule end (*i.e*., both the ‘actin unbinds’ and ‘actin falls behind’ events, n=103). Hence, actin unbinding and actin falling behind the tip are statistically independent of microtubule catastrophes (see supplemental material). Separate channels of the kymographs are shown in Fig. S2. Both TipAct and EB3 are GFP-labelled. Scale bars: (horizontal) and 60 s (vertical).

We observe long-range actin filament transport (Fig. 2F) for actin filaments with a wide range of lengths from around 0.5 μm to 8 μm (Fig. 2G). In the example of Fig. 2A, the actin filament is only 0.6 μm long, which is not much longer than the length of the microtubule tip region where EB3 preferentially binds. Actin filaments that are longer than the tip length sometimes land largely behind the microtubule tip and immediately start following it (Fig. 2B), while others land with part of the filament in front of the tip (Fig. 2C). In the latter situation, we observe that actin transport only starts once the microtubule tip has caught up with the front of the actin filament. Upon analyzing 265 actin transport events, we observe a broad distribution of transport times with a median transport time of 48 s and a long tail that extends up to several minutes (Fig. 2I top, in grey).

The kymographs in Fig. 2A-E demonstrate several mechanisms by which actin transport can end: the actin filament unbinds (21% of events), the microtubule undergoes a catastrophe whereby actin unbinds (48%), the actin filament falls behind the growing microtubule tip region, leaving the filament bound to the microtubule lattice region behind the tip (19%), or the microtubule undergoes a catastrophe while the actin filament remains attached and is pulled along with the shrinking microtubule (11%). We speculate that this backwards pulling of actin could be due to either the presence of residual EB/TipAct on the microtubule and/or actin filaments, or to the peeling protofilaments of the depolymerizing microtubule, where the latter mechanism does not depend on cross-linker binding. CryoEM microscopy of microtubules with end-bound actin might provide more insight into these specific actin transport events, as it allows to study details of the plus-end structure of microtubules [31]. In addition, a small fraction of events (1.5%) corresponds to events truncated by the end of the timelapse movie and one event was ended by a pausing microtubule (shown in Fig. 2E). This pausing event was distinct from regular catastrophe-rescue events, since the location of the microtubule end remained unchanged for a significant amount of time (about 25 s). A summary of the distribution of ending events is shown in Fig. 2H. Irrespective of whether we include all transport events in the distributions or only subsets, the distributions of microtubule growth velocities and of the actin transport times remain similar (Fig. 2I). Specifically, the distribution of transport times excluding those events that end by microtubule catastrophe is indistinguishable from the distribution including all events, in line with the assumption that the three main mechanisms of ending the actin transport, i.e. actin unbinding, microtubule catastrophe, and actin falling behind, are independent Markovian processes (see SI Appendix).

### Simulations Reproduce Microtubule-Mediated Actin Transport

We hypothesize that actin transport can be explained by a competition between two cross-linker induced forces: (i) a driving condensation force that originates from the binding free energy of the cross-linking TipAct proteins that should act to maximize the overlap between the actin filament and the microtubule tip region, and (ii) a friction force due to the presence (at lower affinity) of cross-linkers at the interface between the actin filament and the microtubule lattice that hinders actin transport. To test this hypothesis, we perform kinetic Monte Carlo simulations of the simple model presented in Fig. 3A.

**Figure 3:**
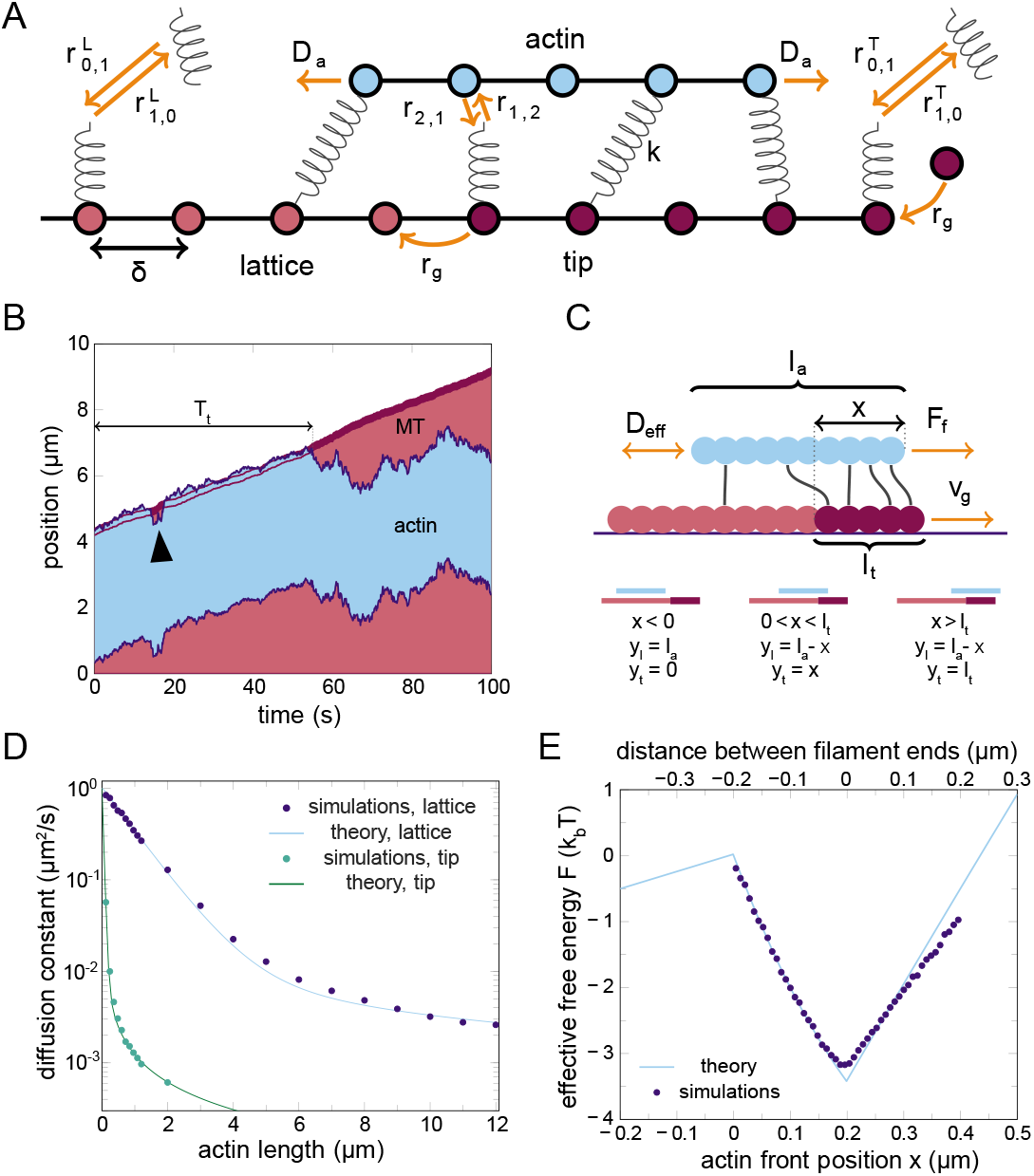
Computer simulations and analytical theory of a mechanism for actin transport by growing microtubules (MTs). (A) The model used for simulating the interaction between an actin filament (blue, top) cross-linked to a growing MT (red, bottom). Both filaments are modeled as one dimensional inflexible chains of binding sites with lattice constant *δ*. The MT grows with rate *r_g_*. Cross-linkers are modeled as springs with spring constant *k* that can be in the solution (state 0), bound to the microtubule only (state 1), or fully connected to both filaments (state 2). These cross-linkers represent a complex of TipAct and EB3, which has a higher affinity for the tip region of the MT (dark red, right) compared to the lattice region (light red, left). The distance between the filaments remains fixed, so the actin filament can only move forward and backward. Viscous interactions with the solution result in a diffusion constant *D_a_* for the actin filament, while the longitudinal components of the pulling forces from the cross-linkers provide additional movement of the actin filament (see also Movie S5). The cross-linker binding rates from solution to the microtubule lattice and tip regions are 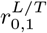, respectively, while the binding rate from a microtubule-bound cross-linker to a fully connected cross-linker is *r*_1,2_. Each transition is microscopically reversible. (B) A typical time trace of the MT and the actin filament. Compare to Fig. 1C. The actin filament is transported when it interacts with the MT tip region (dark red, front of MT), and can recover from quick detachments from this tip region through diffusion (black arrow). However, after a stochastic transport time *T_t_*, the actin filament falls behind the tip region and then performs random diffusion on the MT lattice. (C) Parameter definitions for an analytical theory in a co-moving frame. We define *x* as the position of the front-end of the actin filament compared to the back-end of the MT tip region. Since the tip region advances upon MT growth, this constitutes a co-moving frame of reference. The theory describes the dynamics of *x* using the cross-linker induced effective diffusion constant of the actin filament *D*_eff_ (*x*), the MT growth velocity *v_g_* and the effective forward condensation force *F_f_*. The actin filament has a length *l_a_*, the microtubule tip region has a length *l_t_*, and the microtubule lattice region is assumed to extend leftward. The overlap lengths between the actin filament and the microtubule tip and lattice regions are denoted *y_t_* and *y_ℓ_*, respectively, and the bottom schematics show the relations between these overlap lengths and the other parameters in three regimes. (D) The effective actin diffusion constant *D*_eff_ (*x*) decreases with the overlap between the actin filament and MT lattice region (blue) and the MT tip region (green). Simulations give the proportionality constants of the actin friction coefficients *ζ_t_* and *ζ_ℓ_* by fitting Eq. 1 (lines) to the simulation results (points). (E) Using an analytical expression for the condensation force and the fits from panel D, the theory predicts a free-energy well (blue line), where a co-moving actin position *x* > 0 within the well represents meta-stable transport, whereas a barrier crossing at *x* = 0 and the subsequent slide towards *x* < 0 represents the actin falling behind the MT tip; the free energy is at a minimum at *x* = 0.2 μm where the front of the actin filament is at the front-end of the MT tip. Direct sampling of the positional distribution of *x* in simulations (blue points) confirms the validity of the theoretical prediction. The distance between the two filaments ends (top axis) equals *x* – *l_t_*.

We model the microtubule and the actin filament as one dimensional lattices of binding sites, where the microtubule is immobile while the actin filament can move along its long axis. For simplicity, we assume the same lattice spacing *δ* between the binding sites on both filaments (see SI Appendix). Microtubule growth is modeled through the addition of new binding sites with a fixed rate *r_g_*. We assume that the microtubule contains a chemically different microtubule tip region of *l_t_* = 200 nm long. To maintain a constant size of the tip region, we transform the oldest site in the tip region to a lattice site upon each addition of a new tip site, as indicated in Fig. 3A. We treat TipAct and EB3 proteins as a single complex that can bind to and unbind from the microtubule, justified by fluorescent recovery after photobleaching (FRAP) measurements (Fig. S7B,C). Based on the EB3 fluorescence intensity at the microtubule tip (Fig. S3), we assume that the affinity of the cross-linking complex is 10-fold higher for the tip compared to the lattice region. Since we observe in experiments that TipAct strongly localizes to microtubules, but barely to isolated actin filaments (Fig. 1B), we assume that the cross-linkers always bind to the microtubule first and can then bind to the actin filament. This assumption is further strengthened by the lack of co-localization of EB3/TipAct signal to an actin filament after a transport event, and the high off-rate of TipAct from actin filaments (Fig. S9A). We include the microscopic reverse reactions for all binding reactions, such that microtubule growth is the only process that breaks detailed balance. Free actin filaments diffuse at a rate *D_a_* that depends on the viscosity of the solution. When the actin filament gets cross-linked to the microtubule, its diffusive movement is limited mainly by the transient cross-linker binding events, strongly reducing its effective diffusion constant.

We model the diffusive movement of the actin filament using Brownian Dynamics with a fixed timestep. To allow for movement of the actin filament, the cross-linkers can stretch as Hookean springs with spring constant *k*. Because the cross-linkers are a complex of EB3 and TipAct, and because the spring constant also accounts for the mechanical degrees of freedom of actin filaments, the effective spring constant is lower than observed for other cytoskeletal cross-linking proteins [7, 32–34]. We model the (un)binding of cross-linkers and the addition of new subunits to the microtubule tip as continuous-time Markov processes, which are described by the reaction rates shown in Fig. 3A. We simulate these reactions using a kinetic Monte Carlo algorithm, as described in [7]. We omit microtubule catastrophes from the computational model, since these occur at a constant Markovian rate, as shown in Fig. S1G.

We use experimental observations to estimate all parameters shown in Fig. 3A (for details see SI Appendix). In particular, we measure the actin diffusion coefficient on the microtubule lattice, which depends on the spring constant and binding rates of the cross-linkers. We constrain the spring constant based on the observed actin diffusion constants, and we performed fluorescence recovery after photobleaching (FRAP) experiments on fluorescently labeled TipAct proteins to measure the cross-linker off-rate from microtubules (see SI Appendix and Fig. S7). In addition, we fitted the actin transport time simulated for one specific combination of actin length (4 μm) and growth velocity (3 μm min^-1^) to the order of magnitude of the experimental value of this transport time.

Simulations using this parameter set systematically show that the actin filament follows the growing microtubule tip region, as shown in Fig. 3B. In this example, the actin filament initially overlaps with its front end with the microtubule tip and is transported. At some point it transiently falls behind the tip, but then catches up again (black arrow). We observe the same transient diffusion events in our experiments, as shown in Fig. S4. Eventually the filament completely loses its interaction with the tip and performs random diffusion on the microtubule lattice. Hence, the transition from transport to diffusion occurs when the actin filament loses its interaction with the microtubule tip and does not rebind to the microtubule tip before the microtubule grows away.

### Theory For Cross-linker-Driven Transport And Force Generation

To complement our simulations and identify the requirements for actin transport by growing microtubules, we develop a general coarse-grained theory based on the model schematically shown in Fig. 3C. On timescales longer than the cross-linker (un)binding time, the stochastic motion of the actin filament can be characterised as a biased diffusion process. In this coarse-grained description, the microscopic interactions of the cross-linkers with the filaments give rise to a condensation force that tends to maximize the overlap between the actin filament and the tip of the growing microtubule, and a stochastic force that not only modulates the effective diffusion constant of the actin filament but also generates a friction force that opposes the driving condensation force. We describe this diffusion process via a Fokker-Planck equation, where the actin position *x* is defined in a reference frame that is co-moving with the growing microtubule, as indicated in Fig. 3C. An actin filament moving with the growing microtubule tip thus remains at a constant position *x*.

The actin filament performs 1D diffusion along the microtubule with an effective diffusion constant that depends on the cross-linker binding dynamics. This effective diffusion constant of the actin filament *D*_eff_ decreases with an increasing average number of connections between the actin filament and the microtubule [7, 26, 35, 36]. In turn, the average number of connections depends on the length of the overlap between the actin filament and the microtubule lattice region *y_ℓ_* and between the actin filament and the microtubule tip region *y_t_* (see Fig. 3C). Short actin filaments, or filaments that barely overlap with the microtubule tip, can be in states where no cross-linkers connect the actin to the microtubule. Hence, the effective diffusion constant of the actin filament is the average over the diffusion constants of the unbound and bound states of the actin filament. In the unbound case, the diffusion constant is set by the viscosity of the solution to *D_a_* = 1 μm s ^-1^ (see SI Appendix). In the microtubule-bound state, the actin diffusion constant is dominated by the binding dynamics of the cross-linkers which we capture by an effective friction coefficient *ζ*_bound_(*y_t_*, *y_ℓ_*). The effective diffusion constant is related to this friction coefficient through the Einstein relation, *D*_bound_ = *k_B_T*/*ζ*_bound_ [37]. Put together, the average diffusion coefficient averaged over unbound and bound states is given by

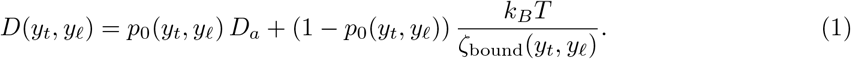

The probability *P*_0_(*y_t_*, *y_ℓ_*) that the actin filament is not bound to the microtubule is given by (see SI Appendix for details)

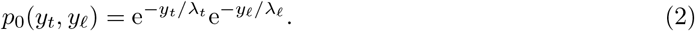

The length scales λ_*t*_ and λ_*ℓ*_ depend on the binding affinities of the cross-linkers for the tip and overlap region, respectively, since these determine the likelihood of cross-linker binding to these regions. We assume that the friction coefficient scales linearly with the actin overlap lengths,

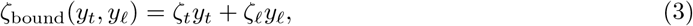

since the average number of cross-linkers is proportional to the overlap lengths *y_t_* and *y_ℓ_*, and friction scales linearly with the number of cross-linkers when the spring constant is low enough and exclusion effects between the cross-linkers are insignificant [26, 35, 36]. To find the proportionality constants *ζ_t_* and *ζ_ℓ_*, we perform two separate sets of simulations where we keep *y_ℓ_* or *y_t_* at 0 and vary the other one, and then measure the diffusion constant of the actin filament. In Fig. 3D, we fit Eq. 1 to the simulated diffusion constants, using *ζ_t_* and *ζ_ℓ_* as the fitting parameter for the curve varying the actin length *y_t_* or *y_ℓ_*, respectively. For small overlap *y*, the effective diffusion constant is dominated by the probability that no cross-linkers are bound, which means that in this regime it decreases exponentially with *y* (Eq. 2), while for large *y* the effective diffusion constant is dominated by the linear increase of the friction coefficient with the number of bound cross-linkers and hence overlap length *y* (Eq. 3). Fig. 3D shows that Eq. 1 fits the diffusion constants in the simulations remarkably well over the whole range of *y*, confirming that the motion of the actin filament is well described by an effective diffusion process.

The diffusive motion of the actin filament is biased forward when the actin filament happens to overlap with the microtubule tip region, because the cross-linkers have a higher affinity for the microtubule tip than for the microtubule lattice. The resulting forward pointing condensation force *F_f_* provides a drift term to this diffusion process. This force is present as long as the actin filament overlaps with the microtubule tip region of size *l_t_*, but does not fully cover it, 0 < *x* < *l_t_*. When the actin filament extends in front of the microtubule tip region, *x* > *l_t_*, a negative, backward condensation force *F_b_* attempts to increase the overlap between the actin filament and the microtubule lattice region. We derive analytical expressions for these forces in the SI Appendix, yielding estimated force values of *F_f_* = 0.10 pN and *F_b_* = −4.6 fN using the parameter set given in Table S1.

Using these forces and Eq. 1, we can describe the effective drift of the actin filament observed in the co-moving frame *x*. The effective net force *F*_eff_(*x*) that acts on the actin filament is the sum of the driving condensation force and the opposing friction force:

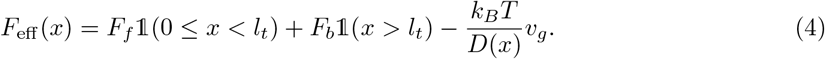

Here, 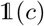 is the indicator function returning 1 if the condition *c* is true and 0 otherwise. The condensation force is thus either the forward driving force *F_f_* when the actin overlaps only partially with the microtubule tip, or the backwards moving force *F_b_* when the actin fully overlaps with the tip but only partially with the microtubule lattice. The diffusion constant *D*(*x*) = *D*(*y_t_*(*x*), *y_ℓ_*(*x*)), where the *x* dependence of the overlap lengths *y* is given in Fig. 3C, and the diffusion constant follows Eq. 1. The last term in the effective force is indeed the opposing friction force, which equals the effective friction coefficient *k_B_T*/*D_x_* times the microtubule growth velocity *v_g_* since that determines the relative speed between the actin filament and the microtubule.

We integrate the effective force from Eq. 4 to find the emergent generalized free energy 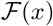 shown in Fig. 3E. The free energy shows three regimes, corresponding to the three situations indicated in the bottom of Fig. 3C. For actin filaments behind the tip (*x* < 0), there is no condensation force since the actin filament does not interact with the microtubule tip region. The negative slope is caused by the effective backward drift of the actin filament in the co-moving frame. For actin filaments that (partially) overlap with the tip (0 ≤ *x* ≤ *l_t_*), the condensation force *F_f_* acts to maximize the overlap region, because the free energy decreases with increasing overlap length. For actin filaments protruding ahead of the tip (*x* > *l_t_*), the overlap between the actin filament and the microtubule tip region can no longer increase, so the forward condensation force disappears. A very small negative backward force *F_b_* remains, since the actin filament loses overlap with the microtubule lattice as *x* increases. However, the largest contribution of the upward slope of the free energy for *x* > *l_t_* is again caused by the effective backward motion in the co-moving frame. The combination of these three regimes creates an energy well, showing that actin transport is a meta-stable state. Interestingly, since the probability distribution *P*(*x*) of the actin position *x* in the simulations is given by *P*(*x*) ∝ exp(−*βF*(*x*)) where *F*(*x*) is the free-energy of the computational model, we can compare the theoretically predicted free-energy profile to −log *P*(*x*) as obtained from the simulations. The generalized free-energy profile obtained from the simulations (symbols in Fig. 3E) closely agrees with the theoretical prediction without any adjustable parameters, supporting the idea that actin transport arises from the balance between a driving condensation force and an opposing friction force.

### Simulations And Theory Correctly Predict Actin Transport Times

The simulations and the analytical theory make predictions about actin transport that can be directly tested against the experimental data. Specifically, we study how the mean duration of transport 〈*T_t_*〉 by growing microtubules changes with the microtubule growth velocity and actin filament length. We observed in Fig. 1C that actin transport ends by microtubule catastrophes, actin unbinding, or actin falling behind the tip region of the microtubule.

The likelihood that cross-linker unbinding will terminate actin transport is low when the overlap between the actin filament and the microtubule tip region is large, *x* ≫ 0, since there will be many cross-linkers holding the actin filament and microtubule tip together. In the other limit that the actin filament falls behind the tip (*x* < 0), the transport event most likely ends by the actin filament rapidly falling even further behind. Hence, the rate at which actin unbinding from the microtubule tip region ends actin transport is dominated by the probability that the actin filament starts to loose overlap with the microtubule tip region, i.e. reaches the top of the free-energy barrier at *x* ≈ 0 (see Fig. 3C). This probability *P*(*x* ≈ 0) can be calculated using the effective free-energy profile, and scales exponentially with the growth velocity *v_g_* (see SI Appendix). Once the actin filament has climbed up the energy barrier for unbinding and lost all connections with the microtubule tip region, unbinding still requires the filament to lose all connections to the microtubule lattice as well. Using Eq. 2, we can calculate how the probability to be disconnected changes with the actin length *l_a_*,

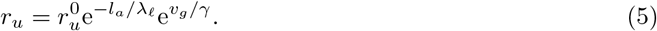

Here, λ_*ℓ*_ is the same as in Eq. 2, and *γ* follows from the free energy and measures how fast the barrier height changes with the microtubule growth velocity *v_g_* (see SI Appendix). Analytical expressions for λ_*ℓ*_ and *γ* are given in the SI Appendix. Only the prefactor 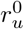 remains unknown, so we fit it once to the complete set of simulated average transport times that is shown in Fig. 4A and B.

**Figure 4:**
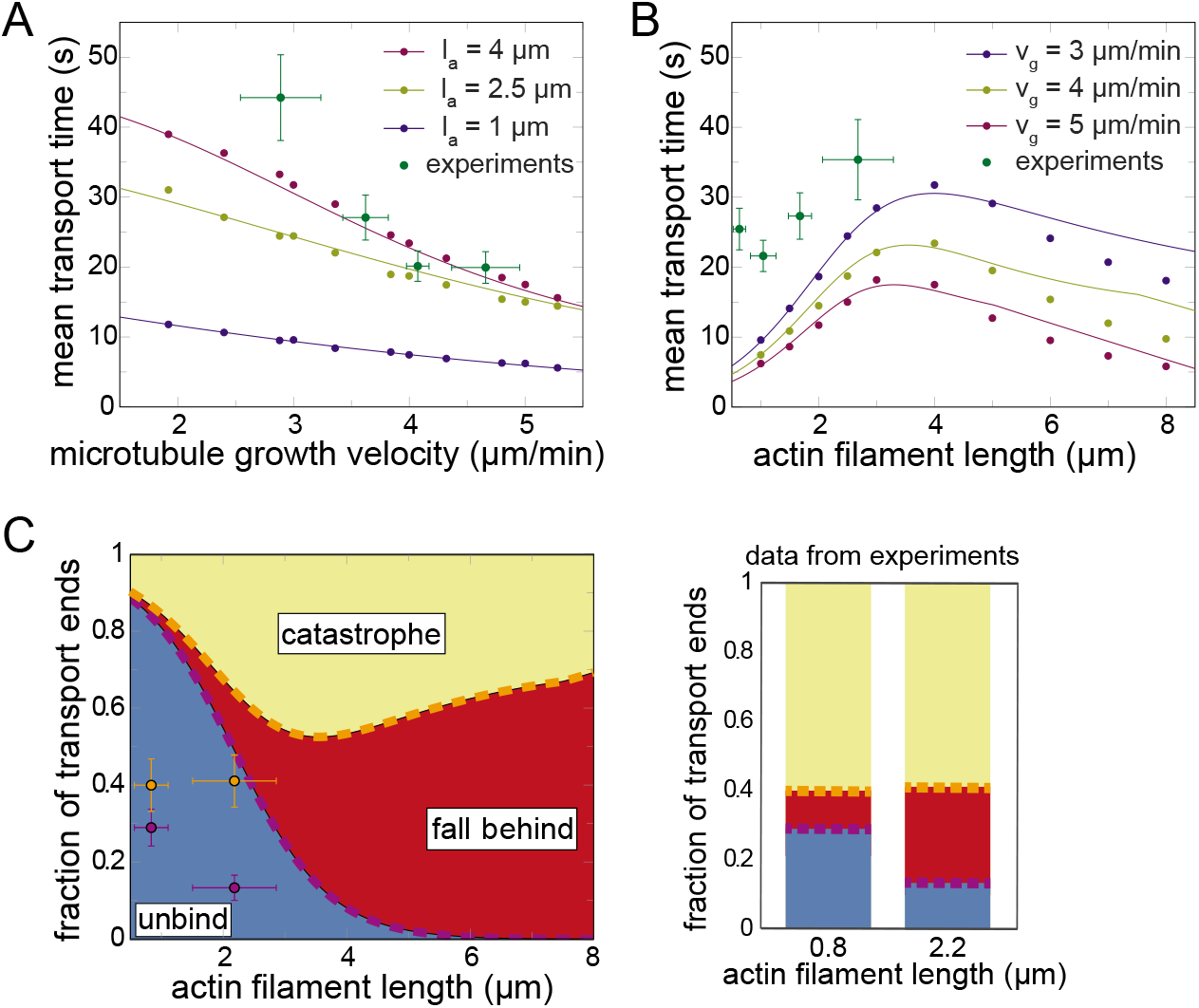
Comparing experimental, simulation, and theoretical results for the mean transport times. The transport time *T_t_* is defined in Fig. 3B. (A) Mean actin transport time plotted against the growth velocity. Simulations and theoretical predictions shown for three different actin filament lengths. Experimental data is sorted according to the growth velocity and grouped into four sets with 63 or 64 data points each. The error bars represent 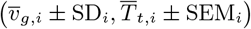, where *i* labels the four groups, a bar on a random variable represents the sample mean, SD is the standard deviation of the distribution of growth velocities, and the SEM is the standard error of the mean of the transport times over that group. The experimental data samples over a distribution of actin lengths 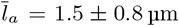 (mean ± SD). The theory correctly predicts both the order of magnitude and the declining trend of the mean transport times. (B) Mean actin transport time plotted against the actin length. Simulations and theoretical predictions shown for three different microtubule growth velocities. The same experimental data as in A is sorted according to the actin length and divided into four equally sized groups of 63 or 64 data points each, with *v_g_* = 3.8 ± 0.7μm min^-1^ (mean ± SD). The experimental data confirms the order of magnitude and the trends of the theoretical predictions. There is insufficient experimental data to test the declining transport times due to increased friction for *l_a_* > 4 μm, and due to increased probability of longer filaments to form bundles, thereby being excluded from transport analysis. However, for *l_a_* < 4 μm, the experimental data confirms that the transport time increases due to an increase of the actin binding times when the actin length increases. (C) Fractions of transport events that end by microtubule catastrophes, actin falling behind the microtubule tip region, or actin unbinding from the tip region as a function of actin filament length. The fractions calculated from theoretical values of the catastrophe rate *r_c_*, rate of falling behind *r_e_*, and unbinding rate *r_u_* are plotted as the indicated colored regions. The indicated data points are calculated from in vitro experiments, and report the borders between the three transport-termination scenarios as shown in barplots on the right. Orange indicates border between catastrophe and fall behind; purple indicates border between fall behind and unbind. We group the experimental data into two bins of equal size (discriminated by actin filament length), because the four data bins used in B provide insufficient statistics per bin to calculate two numbers per bin. We calculate the mean actin length (horizontal error bars are the SD) and the fractions at which each mechanism ends the transport events in both bins. We report the borders between these three regions (vertical error bars are the SEM). Microtubule catastrophes either lead to backward actin transport or to actin unbinding, but are always counted as catastrophes in these fractions. The theory overestimates the fraction of events that unbinds, but it correctly shows that unbinding events become less important with increasing actin length. The overestimation of the unbinding rate also explains the underestimation of the transport times in panels A and B.

To compute the rate at which the actin filament falls behind the tip region, *x* < 0, we note that the filament has to cross a free-energy barrier to leave the tip region, as shown in Fig. 3E. Hence, we can employ Kramers theory [38] to calculate the rate *r_e_* at which the actin filament escapes from the free-energy well,

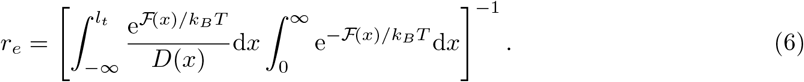

Finally, we treat microtubule catastrophes as an independent process that occurs at a fixed rate *r_c_*. The total rate at which transport ends is given by *r_t_* = *r_u_* + *r_e_* + *r_c_*, and the average transport time is simply the inverse of this rate, 〈*T_t_*〉 = 1/*r_t_*.

We calculate 〈*T_t_*〉 using this theoretical description, and additionally sample *T_t_* in computer simulations as a function of the microtubule growth velocity *v_g_* and the actin length *l_a_*. As shown in Fig. 4, the theory successfully describes the simulation data, showing that the coarse-grained analytical theory is valid for all regimes probed in the simulations. Only the single parameter 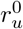 was directly fitted once to the complete set of lines in Fig. 4, otherwise all parameters in the theory are based on analytically calculable expressions or on *ζ_ℓ_* and *ζ_t_*, which are separately fitted to the diffusion constants in Fig. 3D. The curves in Fig.4A show that higher growth velocities lead to shorter transport times. This is because the friction force between the filaments increases while the condensation force remains unchanged, lowering the free-energy barrier for unbinding. Fig. 4B shows that the theory also predicts a decrease of the transport time with increasing actin filament length, again due to an increased friction. However, the transport time is also small for short actin filaments, because in this case transport ends predominantly through actin unbinding. The model therefore predicts an optimal length for processive transport.

Experimentally, we observe a set of transport events, each characterised by a microtubule growth velocity *v_g_*, actin length *l_a_*, and transport time *T_t_*. Since both *v_g_* and *l_a_* vary stochastically in the experiments, we sort the experimental data according to the microtubule growth velocity (or actin length), and group the data into four sets with an equal number of data points each. Then, we calculate the mean 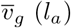 and the mean 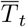 for each set, giving the four green data points in Fig. 4A (Fig. 4B).

The experimental data shows decreasing transport times for increasing growth velocities (Fig. 4A, see also Fig. S6A), a trend that was confirmed by calculations of Spearman correlation coefficients (Table S2). The experiments, having an average actin filament length of 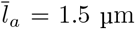, both confirm the predicted theoretical order of magnitude of mean transport times and the trends with varying the growth velocity (Fig.4A). In addition, the experimental data shows increasing transport times for increasing filament length (Fig. 4B, see also Fig. S6B), which was also confirmed by calculations of Spearman correlation coefficients (Table S2). Again, having an average growth velocity of 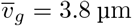, the experimental data confirm the order of magnitude and trends with varying actin length that were predicted by the theory in the small actin length regime (Fig.4B). The experimentally observed actin filaments are not long enough to also probe the regime *l_a_* > 4 μm where the transport time is predicted to decline again due to increased filament friction. The direct unbinding of short actin filaments from the microtubule tip region ends a significant fraction of the transport events, as shown in Fig. 2H. We can also use the theory to predict the fraction of the different events that can terminate transport: microtubule catastrophes, actin falling behind, and actin unbinding. As shown in Fig.4C, we observe that the model parameters overestimate the direct unbinding rate of the actin filaments, but does correctly predict that the fraction of unbinding events decreases strongly with increasing actin filament length.

### Cross-linker Binding Creates A Significant Condensation Force

As explained above, the analytical theory predicts a forward condensation force. To test this prediction, we use optical tweezer measurements in a simplified geometry where TipAct is attached to a bead in the absence of actin filaments (Fig. 5A). Each experiment starts with placing such a TipAct-coated bead in front of a growing microtubule end with the trap. Successful binding of the bead to the microtubule tip region is evident from its movement towards a microtubule (Fig. S9A). This initial movement is a result of the experimental setup and independent of the presence of EB3, as also described before [24]. The bead then gradually returns to the center of the trap as the microtubule grows (Fig. 5A,B), after which it continues to move against an opposing trap force until contact with the microtubule tip is lost and the bead returns to the trap center. We consider the signal starting at the first positive displacement of the bead against the opposing force until the smoothed bead trajectory crosses the zero displacement again as the force signal. Further analysis of these force signals are shown in Fig. S9B. From typical force signals as shown in Fig. 5C, we measure forces with a mean of 0.12 pN (±0.07, SD) for the same conditions as the previously described actin transport experiments (Fig. S9C). The measured force provides an estimate for the condensation force, since the opposing friction force is expected to be very small because of the low velocity.

**Figure 5:**
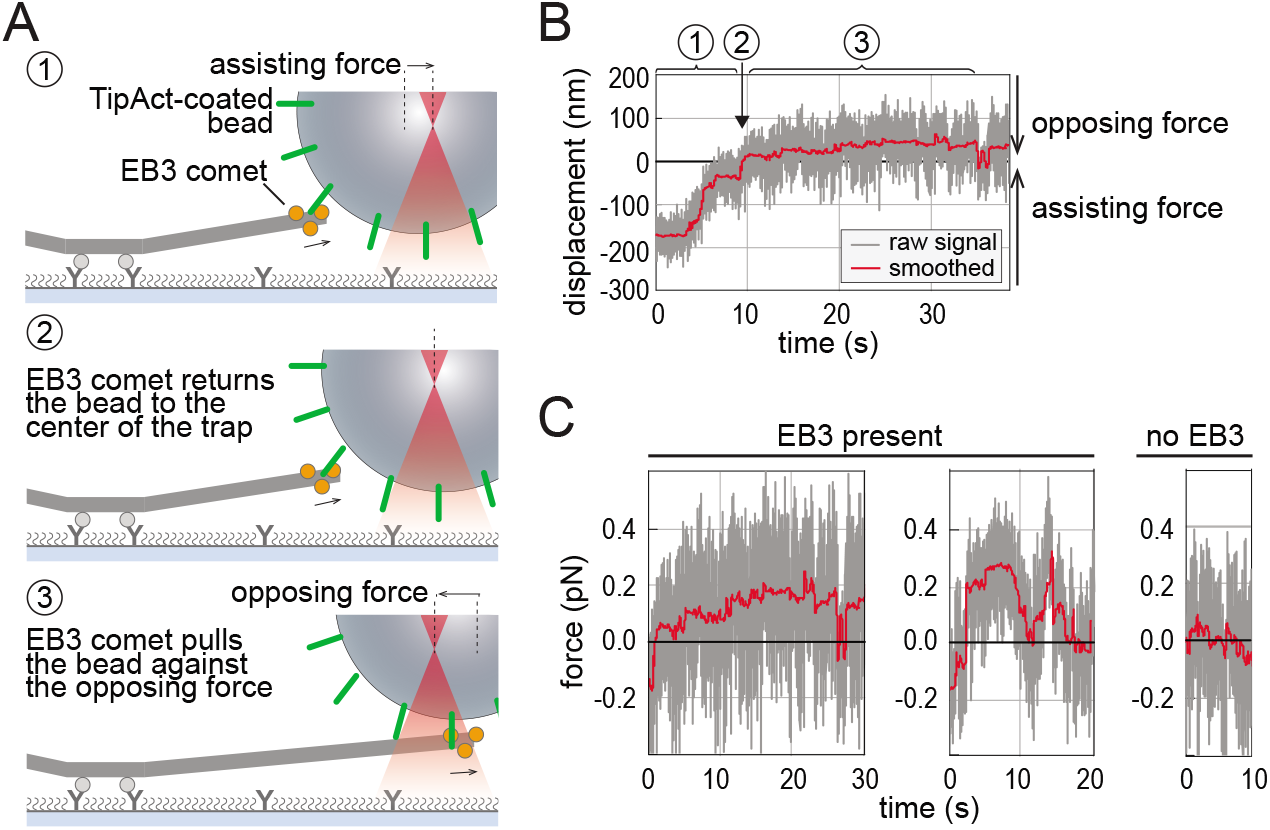
Optical tweezer measurement of the force developed by cargo-bound TipAct at the growing microtubule plus-end. (A) Schematic diagram of the experimental setup. A TipAct-coated bead is initially attached to a growing microtubule carrying an EB3 comet. The bead moves back to the center of the trap under the assisting force (1). After having arrived in the trap center (2), the bead is pulled by the EB3 comet against the opposing force from the trap (3). (B) A typical recording of a bead moving against the opposing trap force with experimental steps numbered according to (A). The large initial movement of the bead is not caused by an active force from the microtubule, but results from an initially free bead that binds to the microtubule. (C). Examples of forces exerted by growing microtubule-ends through the TipAct coupling in presence or absence of EB3. These traces correspond to the complete (un-truncated) force traces in Fig.S9A.

In this simplified experimental geometry, where the TipAct-coated bead replaces the actin filament, our theory predicts a condensation force of roughly 0.2 pN on the bead (Eq. S20 of the SI Appendix), which is close to the value measured experimentally. We also observe experimentally that the force magnitudes decrease when we lower the cross-linker affinity for the microtubule by raising the salt concentration (Fig. S9C), consistent with Eq. S19 of the SI Appendix. This is in agreement with the prediction of our theoretical model, and supports the idea that actin transport is indeed driven by a condensation force.

## Discussion

In this study, we have reconstituted transport of actin filaments by growing microtubule plus-ends in the presence of transiently binding cross-linkers. We have shown that this transport is governed by a condensation force and a competing friction force that are both caused by the cross-linkers. Microtubule growth leads to the net motion of the chemically distinct microtubule tip region. The actin filament recognizes the microtubule tip region through cross-linking proteins that bind more strongly to the tip region compared to the lattice region, creating a force that tries to maximize the overlap between the actin filament and the microtubule tip region. The resulting mechanical movement of the actin filament is opposed by a friction force with the static microtubule lattice, and actin movement is only possible if the cross-linkers adjust their binding positions on the actin filament through (un)binding dynamics. Hence, increasing the condensation force by increasing the cross-linker density does not necessarily lead to longer-range actin transport, since the cross-linker induced friction force between the filaments is also increased.

In recent years, it is becoming increasingly clear that, in addition to motor-driven and (de)polymerization-driven forces, passive cross-linkers can also generate active forces. One mechanism for passive crosslinkers to generate active forces is via entropic expansion [6], where the overlap region between two filaments is maximized to increase the entropy of a fixed number of diffusible cross-linkers, as was shown for Ase1 cross-linkers that connect anti-parallel microtubules in the mitotic spindle [6]. A second mechanism to generate active forces by passive cross-linkers is via their binding affinity for overlap regions when the cross-linkers are present in the solution. This mechanism creates an effective affinity between the filaments, which can result in bundle contractions as was shown for FtsZ filaments in bacteria [25] and for actin filaments [5, 26]. Here we show that this condensation force can also cause directed transport of actin filaments by growing microtubules biased forward by the higher density of cross-linkers present at the growing tip.

The mechanism that we describe is reminiscent of a previously proposed mechanism for chromosome transport via kinetochores, modelled as cargo transport by shrinking microtubules binding in a sleeve [2]. Also in that case, there is a driving force that arises from the affinity of the microtubule for another object (sleeve versus actin filament), and an opposing friction force. However, while in the theoretical analysis of chromosome transport the affinity arises from the interaction between the microtubule and the sleeve, in our model the affinity is mediated by the cross-linkers that prefer to bind from the solution to the overlap region between the microtubule and the actin filaments. Moreover, the former study focused on finding the meta-stable distribution of microtubule positions within the sleeve. In contrast, we not only calculate the effective free-energy profile that emerges from the interplay between the condensation force, the opposing friction force, and the growing microtubule (Fig. 3E), but also the transport time distribution that results from this meta-stable state. Furthermore, while actin transport is driven by the higher affinity of the actin filament for the tip of the growing microtubule than for its lattice, the sleeve-based transport is driven by a depolymerizing microtubule.

Another example of condensation forces that result in processivity is a recent experimental study on pulling of membrane-tubes by growing microtubules [24]. Similar to the transport of actin filaments, the forward force on the membrane tubes is driven by the affinity of the membrane for the growing microtubule plus-end. However, the actin filament slides relative to the microtubule, causing a friction force, whereas the fluid nature of the membrane means that the pulled tubes can spread over the microtubule like a continuous track without creating any friction force.

Within the cell, the condensation force we identified will compete with forces exerted by molecular motors. We showed that actin transport by growing microtubule plus-ends results in much lower forces (0.10 pN) than are measured for microtubule motor proteins (1 pN to 10 pN) [39, 40]. In direct competition, motor forces would therefore likely dominate over the condensation forces. However, condensation forces can remodel actin filaments specifically at the growing microtubule plus-end, whereas motor-driven forces remodel actin filaments along the microtubule lattice, or at depolymerizing microtubule ends [11, 41]. Tip-localized transport of actin filaments may be relevant given recent studies that uncovered two distinct pathways to specifically nucleate new actin filaments at the growing microtubule plus-end [42, 43]. The transport mechanism identified in our study could rapidly relocate those filaments to the leading edge of the cell, where they influence the formation of new actin-based protrusions. In this context, a recent theoretical study explored how such microtubule-based transport could influence the actin distribution within spherical cells [44]. We speculate that growing microtubule plus-ends may also exert significant forces on the cross-linked and branched actin networks they encounter near the cell cortex. Even though cross-linking/branching will prevent actin transport, multiple impinging microtubules could collectively exert a significant force.

The transport mechanism requires transiently binding passive cross-linkers that bind the microtubule end via EB proteins. The most prominent examples of such cross-linkers are the spectraplakins and drebrin [13, 18]. In this work, we used an engineered cross-linker TipAct that contains the actin- and microtubule end-binding domains of ACF7 (also known as MACF1). Full-length ACF7 and its Drosophila homologue Short Stop (also known as Shot) have a more complex functionality because they possess additional domains that bind the microtubule lattice (GAR domain) and regulatory domains of unknown function. It will be interesting to investigate both *in vitro* and in cells how those additional domains modulate the cross-linking and actin transport activity of ACF7. Since actin transport depends on the preferred binding of actin filaments to the growing microtubule-end, we propose to look for these events in cellular contexts where microtubule ends are connected to actin filaments. In mammalian cells, ACF7-mediated actin-microtubule interactions take place at the ends of microtubules that impinge on the protrusive front of the cell [10]. In Drosophila, Shot-mediated interactions were recently identified at the ring canals of the ovary, where actin-microtubule cross-linking is necessary to control the transport of cytoplasmic materials from nurse cells to the oocyte [45]. A similar transport mechanism may also contribute to the transport of intermediate filaments by microtubules [46–49].

In our study, the actin filaments were stabilized so that their length is essentially constant as a function of time. Our model predicts that the transport of these filaments is accompanied by a steady-state overlap length, which arises from the balance between the driving condensation force, which is independent of the overlap length and the total actin length, and the opposing friction force, which increases linearly with not only the overlap length but also the total actin length (see Fig. 3E). In vivo, actin filaments are however dynamic: they can grow, shrink, or even treadmill, in a direction that depends on their polarity. All three processes can change the overlap length and/or the total actin length, and thereby affect the balance between the driving condensation force and the opposing friction force. We anticipate that actin growth would impede transport by increasing the friction. In contrast, treadmilling does not change the total length. It will therefore not significantly affect transport when it is slow compared to the transport speed as set by the microtubule growth rate, because in this quasi-equilibrium regime the overlap rapidly adjusts to the treadmilling to maintain the steady-state overlap length. Yet, when treadmilling is faster, it is likely to enhance transport when it occurs in the direction of microtubule growth. We leave a quantitative analysis of these regimes for future work. Finally, the theoretical description we present may be applicable not only to filament transport, but also to transport by propagating reaction-diffusion patterns on membranes, such as the transport of plasmids by ParA [50–52].

## Material and methods

### Protein Isolation and Preparation

Engineered cytolinking protein TipAct (composed of an N-terminal eGFP followed by the actin binding domain and EB-binding domain of full-length MACF1, separated by the coiled-coiled linker of Cortexillin I) was expressed in E. coli T7 cells and purified using a His-tag affinity column, as was previously described [15, 53]. Lyophilized porcine brain tubulin was obtained from Cytoskeleton (Denver, CO, USA), resuspended at 50–100 μM in MRB80 buffer [80 mM piperazine-*N*,*N*’-bis(2-ethanesulfonic acid) (Pipes), pH 6.8 with KOH, supplemented with 4 mM MgCl_2_ and 1 mM EGTA], snap-frozen and stored at −80°C until use. G-actin was purified from rabbit skeletal muscle acetone powder as previously described [54], filtered on a Superdex 200 10/60 size-exclusion column (GE Healthcare, Waukesha, WI, USA) to remove any oligomers and stored at −80°C in G-buffer [5 mM tris(hydroxymethyl)aminomethane (Tris), pH 7.8 with HCl, 0.1 mM CaCl_2_, 0.2 mM ATP, 1 mM dithiothreitol (DTT)]. Fluorescent actin was prepared by labelling monomers with Alexa Fluor 649 carboxylic acid succinimidyl esther (Molecular Probes, Life Technologies, Carlsbad, CA, USA). Before use, G-actin was thawed overnight at 4°C and spun for 15 min at 149,000 × g to remove any aggregates, and stored at 4°C for no longer than 2 weeks. 6xHis-tagged recombinant human GFP-EB3 and mCherry-EB3 were expressed and purified as described before [55].

### Dynamic Microtubule Assay

Microscope flow cells were constructed as previously described [53]. In brief, flow channels of 10-15 μl, assembled from base-piranha cleaned glass coverslips and slides and Parafilm spacers, were functionalized by sequential incubation with 0.1 mg/ml PLL-PEG-Biotin (PLL(20)-g[3.5]-PEG(2)/PEG(3.4)-Biotin(20%), SuSos AG, Dübendorf, Switzerland) for 30–60 min, 0.25 mg/ml streptavidin (Thermo Scientific Pierce Protein Biology Products, Rockford, IL, USA) for 10 min, 0.5 mg/ml κ-casein for 10 min and 1 % (w/v) Pluronic F-127 for 10 min, all solutions in MRB80, and including 40–70 μl rinses with MRB80 in between incubation steps.

Actin filaments were pre-polymerized at 7.5 μM G-actin concentration (15 mol % labelled and 85 mol % unlabelled premixed monomers) in MRB80 for 30–90 min at room temperature, before adding phalloidin (Sigma-Aldrich) in 1:1 molar ratio to stabilize the filaments. Dynamic microtubules were nucleated from guanylyl-(α, β)-methylene-diphosphonate (GMPCPP) stabilized microtubule seeds, which were bound to the biotinylated surface and prepared according to standard double-cycling protocols [53,56]. Any non-attached seeds were rinsed off with MRB80 before adding the final imaging mix. The core imaging buffer consisted of MRB80 supplemented with 0.5 mg/ml κ-casein, 0.2% (v/v) methyl cellulose (M0512, Sigma-Aldrich Chemie B.V., the Netherlands), 75 mM KCl, 1 mM guanosine triphosphate(GTP), 0.2 mM Mg-ATP and an oxygen scavenging system [4 mM dithiothreitol (DTT), 2 mM protocatechuic acid (PCA) and 100 nM protocatechuate-3,4-dioxygenase (PCD)]. This solution futher contained tubulin concentrations ranging from 20 to 30 μM (6 mol % Rhodamine labeled), 7.5-10 nM F-actin, 133 nM EB3 (mCherry or GFP labeled), and 50 nM TipAct (GFP).

### Microscopy

Imaging of *in vitro* actin transport by growing microtubule plus-ends was performed using a Nikon Eclipse Ti-E inverted microscope (Nikon Corporation, Tokyo, Japan) equipped with an Apo TIRF 100x/1.49 numerical aperture oil objective, a QuantEM:512SC EMCCD camera (Photometrics, Roper Scientific), a motorized stage, Perfect Focus System, and a motorized TIRF illuminator (Roper Scientific, Tucson, AZ, USA). For excitation, we used a 561 nm (50 mW) Jive (Cobolt, Solna, Sweden), a 488 nm (40 mW) Calypso (Cobolt) diode-pumped solid-state laser and a 635 nm (28 mW) Melles Griot laser (CVI Laser Optics & Melles Griot, Didam, Netherlands). A custom-built objective heater was used to maintain the sample at 30°C. Triple-colour images were acquired for 10-20 min with 3-s intervals and 100-200 ms exposure times.

### Image and Data Analysis

Image processing and analysis were performed using plugins for Fiji [57] or ImageJ and custom-written programs in Python or MATLAB. Timelapse series of TIRF images were drift-corrected using a custom-written Matlab program. Kymographs of microtubule growth were created using the reslice tool in Fiji [57]. Plus- and minus-ends were distinguished based on their growth rates: faster-growing ends were identified as the plus-ends and the slower-growing ends as the minus-ends. The parameters characterizing actin transport and microtubule dynamics were determined from these timelapses and kymographs.

A microtubule/actin interaction event was classified as transport and included in further analysis when (1) an actin filament colocalized with an EB3/TipAct-complex at the growing microtubule plus-end, (2) moved for at least 0.5 μm (3 pixels) and for 9 sec (3 pixels) along with the microtubule tip, and (3) was not interacting with other actin filaments or microtubules. The actin filament length and tracking time were obtained from manual fits on the kymographs. From the original time lapse image series, the binding states of the actin filament before and after the transport event were determined.

The microtubule dynamics were characterized as previously described [58]. In brief, growth velocities were obtained from manual fits to the growth phases, and the average velocity was taken as the average over all events weighted with the duration of the individual events. The error is the weighted standard deviation. Catastrophe rates were calculated as the number of catastrophe events divided by the total time microtubules spent growing. The error is given by the rate divided by the square-root of the number of events.

### Simulations and theoretical modeling

The simulations are based on a modified version of the algorithm described in [7]. All details are described in the SI Appendix.

### Optical Tweezers

Silica beads (1 μm in diameter) functionalized with carboxy groups (Bangs Laboratories) were conjugated PLL-PEG-Biotin (PLL(20)-g[3.5]-PEG(2)/PEG(3.4)-Biotin(10%), SuSoS AG, Switzerland) as described [59]. 10 μL of PLL-PEG-coated beads (ca 0.2% w/v) were washed in washing buffer (80mM K-Pipes pH 6.9 with 1mM EGTA, 4mM MgCl_2_, 1mM DTT and 0.5 mg/ml casein) by centrifugation for 1 min at 16000 g, then resuspended in 20 μL of 8 μM Neutravidin (Thermo Scientific) in the same buffer. After 30min incubation at 23°C with frequent mixing, the beads were washed twice in 100 μL of the washing buffer, resuspended in 20 μL of 0.2 μM biotinylated anti-GFP IgG (Rockland) and further incubated for 30 min at 23°C with frequent mixing. After two more washes in the washing buffer, the beads were resuspended in 20 μL 120nM GFP-TipAct and incubated for 1 hour on ice with frequent mixing. Finally, the beads were washed 3 times and resuspended in 50 μL of the washing buffer.

To attach microtubule seeds to glass, we used digoxigenin (DIG)-labelled tubulin to prevent interactions with the biotinylated beads. Briefly, tubulin was purified from porcine brain [60] and chemically labelled with NHS-DIG [61]. GMPCPP stabilized microtubule seeds were prepared using 30% DIG-labelled tubulin and according standard double-cycling protocols [56].

Optical trapping was performed as described previously [24]. Slides and coverslips were silanized using Plus-One Repel Silane (GE Life Sciences), then assembled into flow chambers using double-sided tape and functionalized with 0.2 μM anti-DIG IgG (Roche), then passivated with 1% Pluronic F-127 in MRB80, before DIG-labelled seeds were introduced. The reaction mix (80 mM K-Pipes pH 6.9, 50 or 75 mM KCl, 1 mM EGTA, 4 mM MgCl_2_, 1mM GTP, 1 mg/ml κ-casein, 4 mM DTT, 0.2 mg/ml catalase, 0.4 mg/ml glucose oxidase, 20 mM glucose and either 25 mM unlabelled tubulin with addition of 133 nM EB3, or 10 mM unlabelled tubulin in the absence of EB3) was centrifuged at 30 psi for 5 min. TipAct-coated beads were added to the supernatant at a ratio of 1:33-1:50. This final mix was then introduced into the flow chamber, and experiments were carried out at 25°C.

DIC microscopy and optical trapping were performed as described previously [59]. Measurements were performed at nominal trap power of 0.2W (the lowest setting of our instrument), and then the trap stiffness was additionally softened down to 4-6·10-3 pN/nm using a circular polarizing filter placed in the wave path of the trapping laser. The QPD signal was recorded at 10 kHz.

## Data Availability

Additional data are available in the SI Appendix. Original data from *in vitro* actin transport experiments, optical tweezer measurements and simulations will be available via https://doi.org/10.4121/c.5584335

## Acknowledgements

We thank Jeffrey den Haan and Marjolein Vinkenoog for actin purification, and Andrea Martorana for TipAct purification. We thank Michel Steinmetz for the gift of EB3’s. We thank Maurits Kok for help with image analysis. We thank Michele Monti for many useful discussions. This work was supported by the European Research Council (Synergy grant 609822 to M.D. and A.A.).

## Author contributions

Author contributions: C.A., H.W., V.A.V., P.R.t.W., M.D. and G.H.K. designed research; C.A., H.W., V.A.V. and M.P.L. performed research; C.A., H.W. and V.A.V. analyzed data; M.P.L and A.A. edited paper; and C.A., H.W., V.A.V., P.R.t.W., M.D. and G.H.K. wrote the paper.

The authors declare no competing interest.

## Notes

### Competing Interest Statement

The authors have declared no competing interest.

### Summary of Updates

Figures 2A, 2E, and 4C revised. Discussion section updated with biological relevance and with implications dynamic actin filaments. Supplemental files updated. Minor typos corrected and clarifications added.

