## Supplemental Information for "Cross-linkers at growing microtubule ends generate forces that drive actin transport"


### Binding kinetics of actin transport

As input parameters for the simulations and theoretical model, we measured the relevant binding kinetics, as shown in Fig. 7. This section discusses the methods and results.

To test the binding affinity of TipAct to actin filaments, we measured single-molecule dwell times of TipAct on actin filaments (Fig. 7A). Phalloidin-stabilized actin filaments were bound to the coverslip surface via biotin-streptavidin links. Thereafter, TipAct was added to the flow cell at a concentration of 25 nM and TIRF microscopy imaging was performed at 33 ms/frame. We observed that TipAct barely binds to actin filaments when no microtubule is present, and the binding events that do happen are quickly followed by unbinding events. Fitting a single-exponential to the normalized distribution of TipAct dwell times (Fig. 7A, right) yielded an average dwell time of $(111 \pm 6)$ ms (n=301) and corresponds to a single-molecule off-rate of about 9 $s^{-1}$.

To probe protein off-rates at microtubule plus-ends, we performed fluorescence recovery after photobleaching (FRAP) measurements (Fig. 7B,C). We analyzed the recovery curves for mCherry-EB3 and GFP-TipAct at both free and actin-bound microtubule tips. After normalization, we fitted the data to estimate the unbinding rates. For the free microtubule tips (i.e. not bound to an actin bundle), a reaction-limited recovery curve multiplied by a decaying single-rate exponential envelope was sufficient to describe the recovery. Thus, the equation used to fit the free MT tips data was: $I(t) = (A - B * exp[-t * k_{off-free}]) * exp[-H(t - t_0) * (t - t_0) * k_{decay-free}] + D + (1 - exp[-H(t - t_0) * (t - t_0) * k_{decay-free}]$, where A, B and D are constants, $k_{off-free}$ the off-rate of mCherry-EB3 (or GFP-TipAct) at microtubule tips, $k_{decay_{free}}$ the transition rate from a tip-like to a lattice-like binding profile (related to GTP hydrolysis), and $H(t - t_0)$ the Heaviside function to account for the fact that this transition occurs with a time delay. For microtubule tips bound to actin, we used a modified version of this equation to fit the data, namely: $I(t) = (A - B * exp[-t * k_{off-free}] - C * exp[-t * K_{off-bound}]) * exp[-H(t-t_0) * (t-t_0) * k_{decay-bound}] + D + (1 - exp[-H(t-t_0) * (t-t_0) * k_{decay-bound}]$, where $k_{off-free}$ is the off-rate of mCherry-EB3 (or GFP-TipAct) bound only to microtubule tips and $k_{off-free}$ is the off-rate of mCherry-EB3 (or GFP-TipAct) bound both to the microtubule tip and actin.

After fitting the recovery curves to these equations, we found unbinding rates from free microtubule tips of $3.1\,s^{-1}$ and $1.8\,s^{-1}$ for TipAct and EB3, respectively (Fig. 7C). For actin-bound microtubule tips, we found unbinding rates of $0.7\,s^{-1}$ and $0.8\,s^{-1}$ for TipAct and EB3, respectively (Fig. 7C). The unbinding rates from actin-bound microtubule tips are comparable for TipAct and EB3, indicating that we can consider them as a single EB3/TipAct-complex.

### Effect of microtubule catastrophes on the transport time distribution

As shown in Fig. 2H of the main text, transport events typically end in one of three ways: (i) actin can fall behind the growing microtubule tip and escape to the microtubule lattice, (ii) actin can unbind directly from the microtubule tip and escape to the solution, or (iii) a microtubule catastrophe occurs. We call stochastic times at which these events happen $T_e$, $T_u$, and $T_c$, respectively. Our model assumes that these three random variables are independent, and the shortest time decides which event actually occurs. In this section, we discuss a framework to separate the events that end with the actin falling behind the tip or unbinding from simple catastrophe events. We are interested in this separation of events, because, accounting for the cross-linker properties in our model, we want to check whether it can explain how long actin filaments remain associated with growing microtubule tips. Hence, we group the former two event types, $T_i \equiv \min(T_e, T_u)$, and refer to them as events of interest. In case of a microtubule catastrophe, the actin filament either unbinds from the microtubule or it is transported back by the shrinking microtubule (see Fig. 2B and Fig. 2D), but we consider the microtubule catastrophe itself to be responsible for terminating transport in both cases, since we only focus on forward transport. We call $r_i = r_e + r_u$ the rate at which the events of interest occur, and we call the rate of microtubule catastrophes $r_c$. Then, the experimentally measurable rate $r_t$ at which the transport events end is given by

$$r_t = r_i + r_c. \tag{1}$$

The observed transport time $T_t$ obeys

$$T_t = \min(T_i, T_c)\,. \tag{2}$$

Since events ending with versus without a microtubule catastrophe are independent and exponentially distributed, we have

$$\mathbb{P}(T_t > t) = \mathbb{P}(T_i > t \wedge T_c > t) = \exp[-\,(r_i + r_c)\,t]\,, \tag{3}$$

consistent with Eq. 1.

If we cut out the events that end in catastrophes, we observe the transport time distribution

$$\mathbb{P}(T_t > t \mid T_i < T_c) = \frac{\mathbb{P}(T_i > t \wedge T_i < T_c)}{\mathbb{P}(T_i < T_c)}, \tag{4}$$

where we rewrite the first probability using Bayes' theorem and Eq. 2. We can calculate the probability density of $T_c$ (events ending by a microtubule catastrophe) happening at time $t'$, and integrate over all possible $t'$,

$$\begin{aligned}
\mathbb{P}(T_i > t \wedge T_i < T_c) &= \int_0^\infty \mathbb{P}(t < T_i < t')\, r_c \exp[-r_c t']\,\mathrm{d}t' \\
&= \int_t^\infty (\exp[-r_i t] - \exp[-r_i t'])\, r_c \exp[-r_c t']\,\mathrm{d}t' \\
&= \frac{r_i}{r_i + r_c} \exp[-\,(r_i + r_c)\,t]\,. 
\end{aligned} \tag{5}$$

In a similar fashion, one can show that

$$\mathbb{P}(T_i < T_c) = \frac{r_i}{r_i + r_c}. \tag{6}$$

Hence, Eq. 4 gives

$$\mathbb{P}(T_t > t \mid T_i < T_c) = \exp[-\,(r_i + r_c)\,t] = \exp[-r_t t] = \mathbb{P}(T_t > t)\,. \tag{7}$$

The cumulative distribution function (CDF) we look for is simply $\mathbb{P}(T_t \leq t \mid T_i < T_c) = 1-\mathbb{P}(T_t > t \mid T_i < T_c)$, and Eq. 7 shows that this distribution equals the CDF $\mathbb{P}(T_t \leq t)$. Naively, one may expect that removing the catastrophe data and looking only at the events of interest would lead to measuring the CDF $\mathbb{P}(T_i \leq t)$, the distribution that is measured when catastrophes never occur. However, the actual distribution in Eq. 7 is biased compared to $\exp[-r_i t]$. To explain this, it is helpful to estimate the likelihood that an event ends in a catastrophe when the event time $T_i$ is short ($T_i < 1/r_c$) or when it is long ($T_i > 1/r_c$). When $T_i$ is short, and catastrophes do occur, it is likely that the catastrophe takes place after the actin filament escapes, or $T_i < T_c$. When the events that end in catastrophe are then discarded, it is likely that the short $T_i$ is still observed. However, when $T_i$ is long, the catastrophe is more likely to happen before the event of interest, and usually $T_i > T_c$. The selection then discards these events more often. Hence, short $T_i$ are selected more often than long $T_i$, which introduces a negative bias in the distribution of transport times of those events that do not end in a microtubule catastrophe. Eq. 7 shows that the CDF for the selected transport times $\mathbb{P}(T_t \leq t \mid T_i < T_c)$ equals the CDF of the complete data set of transport times $\mathbb{P}(T_t \leq t)$, and is biased compared to the desired CDF $\mathbb{P}(T_i \leq t)$. Hence, we proceed by simply measuring the mean transport time using all events $T_t$, and then separately measure the microtubule catastrophe rate in order to separate the contributions of $r_i$ and $r_t$.

### Cross-linker parameters in the simulated model

The simulation model (see Fig. 3A of the main text) is based on a model previously developed to describe microtubules [1] interacting via cross-linker proteins. It uses discrete time step dynamics for the movement of the actin filament, while using a kinetic Monte Carlo algorithm for simulating the Markovian reactions that take place with time-varying rates. Here we explain in detail how the actin-microtubule cross-linkers are implemented in the simulation. The parameters are based on the experimental characterization of the interactions of TipAct with actin filaments and microtubules. TipAct has a binding site that can bind to actin directly [2]. In the presence of EB3, we observe that TipAct has a higher binding affinity for microtubules than for actin filaments, as is indicated by the co-localization of TipAct to growing microtubule ends as opposed to co-localization of TipAct to actin filaments (Fig. 1B of the main text, quantified in Fig. 7). TipAct interacts with microtubules through EB3, which in turn has a high affinity for the microtubule tip region and a low affinity for the microtubule lattice region [3]. In the model, we make the simplifying assumption that there is a single cross-linker that represents the complex of TipAct and EB3, but could also represent states where TipAct binds to microtubules and actin on its own. We will refer to this single cross-linker as TipAct, except when we need to specifically distinguish between internal binding states.

To form a cross-link between a microtubule and an actin filament, there are two different pathways. TipAct can first bind to the microtubule or first to the actin filament. Next, it can bind to the opposite filament. All these binding transitions are reversible. Hence, there are eight (un)binding rates; two for the transition between the unbound and actin-bound states, two between the unbound and microtubule-bound states, two between the actin-bound and fully bound states, and two between the microtubule-bound and fully bound states. Detailed balance reduces this to a set of seven different rates. Our *in vitro* experiments show that the binding of TipAct to single actin filaments is negligible compared to the binding to single microtubules (Fig. 1B, Fig. 1C and Fig. 7), meaning that we can remove one of the pathways, and we are left with four (un)binding rates per pair of binding sites on the microtubule and actin filament, as shown in Fig. 3A of the main text. These are the two rates for the transition between a fully unbound state and a state where TipAct is solely bound to the microtubule, and two rates for the transition from this dangling cross-linker state to the state where the cross-linker binds both the microtubule and the actin filament.

Since TipAct binds strongly to the microtubule tip region through EB3, but it also has a low but significant affinity for the microtubule lattice through EB3, we use two separate sets of (un)binding rates for TipAct binding from the solution to the microtubule lattice and tip regions. However, we assume that the rates to bind to or unbind from the actin filament are the same for dangling TipAct cross-linkers bound to the microtubule tip or lattice. For convenience, we label the unbound state as 0, the state with a dangling cross-linker bound to the microtubule as 1, and the fully bound state where the cross-linker also binds to the actin filament as 2, and we distinguish between the microtubule lattice region $L$ and the microtubule tip region $T$. As Fig. 3A shows, this gives us the rates $r^T_{0,1}$, $r^T_{1,0}$, $r^L_{0,1}$, $r^L_{1,0}$, $r^0_{1,2}$, and $r^0_{2,1}$. Here, a rate $r_{i,j}$ corresponds to the transition from state $i$ to state $j$, and the 0 in the superscript indicates that the rate will depend on the amount that the linker has to stretch,

$$r_{1,2}(d) = r^0_{1,2} \exp\left[-\frac{U(d)}{2k_BT}\right], \tag{8}$$

where $d$ is the amount by which the linker is stretched after the transition, and $U(d)$ gives the potential energy for a cross-linker with extension $d$. The same equation holds for the reverse rate $r_{2,1}(d)$ if we change the sign of the potential energy. Detailed balance sets a Boltzmann factor for the ratio between the forward and backward rates, but not for the rates individually. We make the model choice to distribute this factor evenly between the forward and backward reactions, explaining the factor $1/2$ in the exponent in Eq. 8. This choice ensures that the binding rate to a site that requires a cross-linker stretch $d$ decreases with $d$, and the corresponding unbinding rate increases with $d$.

Even though there are typically multiple cross-linkers connecting the actin filament to the microtubule, the filaments can occasionally disconnect due to the stochastic nature of the (un)binding dynamics of the cross-linkers. However, upon the unbinding of the last remaining cross-linker, the filaments are still in close enough proximity such that new cross-linkers can bind. Hence, cross-linker rebinding can keep the filaments connected. To simulate this rebinding dynamics, we introduce a time scale $\tau_a$ and end a simulation when there are no connections between the two filaments for longer than $\tau_a$. This time scale accounts the time it takes the actin filament to diffuse over a distance corresponding roughly to the size of a cross-linker, such that the probability of rebinding becomes negligible [4], and is estimated in Eq. 11.

Since we keep the distance between the filaments fixed in the simulations, we only have to describe the longitudinal component of the stretch of the cross-linkers $d$, and we assume that the potential energy obeys a simple harmonic function $U(d) = kd^2/2$. This potential provides forces on the actin filament and also influences the (un)binding rates through Eq. 8. When a cross-linker is bound to the microtubule and dangling, there are many binding sites on the actin filament that the cross-linker could bind to, but the harmonic potential sets very low rates for binding to the ones that require the cross-linker to stretch far. Hence, we improve the efficiency of the simulations by setting a maximum stretch for the cross-linkers, such that the simulations do not check binding positions that are unrealistically far away. We set this maximum stretch to $4\delta$, where $\delta$ is the spacing between binding sites on either filament. Using the parameter values from Table 1, the springs store a potential energy of $10.4\,\mathrm{k_BT}$ at a stretch of $4\delta$. Hence, thermal fluctuations are unlikely to cause a stretch that large, justifying the simulation choice. For consistency, we also disallow the actin filament to change its position in a way that would stretch linkers more than $4\delta$.

Another model choice we explain in the main text is that the density of microtubule tip sites follows a step function, such that the tip is a well-defined region on the microtubule of length $l_t$. This simple structure simplifies calculations and expresses our ignorance about the precise structure of the microtubule tip region where EB3 binds, which remains under discussion [3, 5, 6]. To confirm that the proposed mechanism of tip tracking also works with different microtubule tip region structures, we performed simulations where the density of tip sites decreases exponentially with the distance from the front of the microtubule. Leaving the other parameter values as they are listed in Table 1, we still observed actin transport, as shown in Fig. 10. Hence, the mechanism of a forward condensation force caused by transiently binding cross-linkers works independent of the exact shape of the tip region.

### Actin filament diffusion measurements

We experimentally observed that actin filaments occasionally land on the microtubule lattice and perform one-dimensional diffusion while remaining bound via TipAct. By raising the tubulin concentration to 30 µM, we obtained longer microtubules that allowed us to measure the diffusive motion of the actin filaments on the microtubule lattice, away from the tip region. The estimation of the diffusion constant is complicated by thermal fluctuations of the apparent actin filament length as observed by TIRF illumination. We quantify this apparent length by manually measuring the positions of the beginning and the end of the actin filament, called $x_b$ and $x_e$, respectively, as a function of time with time steps of $\delta t = 3\,\mathrm{s}$. Two kymographs of diffusing actin filaments are shown in Fig. 5AB, along with the trajectories of $x_b$ and $x_e$ and the apparent actin length $l_a = x_e - x_b$. Besides small fluctuations around a constant length, we observe large transient deviations of the actin length at the beginning of the time traces (in the bottom panels of Fig. 5AB). These length fluctuations are probably caused by the binding process of the actin filaments, since binding will bring the whole actin filament from solution closer to the surface, therefore bringing it in the TIRF field. To estimate the length of the part of the actin filament that interacts with the microtubule, we remove the initial part of the time traces and take the mean value of $l_a$ over the cropped time trace, as shown in Fig. 5AB. Then, we define the actin position $x$ as the center point of the actin filament, $x = (x_b + x_e)/2$, and estimate the diffusion constant from the time trace of $x$ in the cropped window.

Since the recorded data for $x$ is noisy and contains a finite number of time points, it is difficult to find a reliable estimate of the diffusion constant from a single time trace [7]. The data set consists of $N + 1$ time points $x_i$, which are acquired at $t_i = i\delta t$ and $i = \{0, 1, \ldots, N\}$. For the diffusion constant, we use a family of estimators based on the time-averaged square displacement [8],

$$\overline{D}_\Delta = \frac{1}{2\Delta\,(N - \Delta + 1)\,\delta t} \sum_{i=0}^{N-\Delta} \left(x_{i+\Delta} - x_i\right)^2 , \tag{9}$$

which depends on a choice of the number of time steps $\Delta \in \mathbb{N}$ over which we probe the fluctuations. The movement of the actin filament is likely correlated for small times, and when $\Delta$ is smaller than the largest correlation time, the estimate of the diffusion constant will be influenced by these correlations. Furthermore, measurement noise and thermal length fluctuations have a larger influence on $\overline{D}_\Delta$ when $\Delta$ is small, because the actin steps $(x_{i+\Delta} - x_i)$ are typically smaller. However, when $\Delta$ is too large, the number of time points $N - \Delta$ decreases, and the uncertainty in the estimate grows. As shown in in Fig. 5D, we measured time traces that last for only 30 s to 950 s before cropping, giving between 9 and 291 data points after cropping. We focus on the two time traces that are longer than 500 s to obtain reliable estimates of the actin diffusion constants.

Since estimates of the diffusion constant will be influenced by the number of time steps, we show the results of Eq. 9 as a function of the time gap $\Delta\delta t$ for the two long time traces in Fig. 5C. These two time traces are obtained from actin filaments with lengths $l_a = 6.6\,\mu\mathrm{m}$ and $l_a = 7.1\,\mu\mathrm{m}$, and after cropping the time traces to a total duration of 873 s and 762 s, respectively. For small times, the estimate $\overline{D}_\Delta$ decreases strongly with $\Delta$ and reaches a plateau value after roughly 150 s. We believe that the strong bias in $\overline{D}_\Delta$ contains contributions from actin filament length fluctuations and external measurement noise. Furthermore, actin movement is different for time scales shorter than the time scales over which cross-linker remodeling occurs, and we expect that these effects also contribute to the bias in $\overline{D}_\Delta$. We therefore estimate the true diffusion constant as the sample mean of $\overline{D}_\Delta$ over a window between 150 s and 450 s, and report the standard deviation of the mean over this window as the error in our estimate. This results in the values $\overline{D} = 0.014(3)\,\mu\mathrm{m}^2\,\mathrm{s}^{-1}$ ($l_a = 6.6\,\mu\mathrm{m}$) and $\overline{D} = 0.009(3)\,\mu\mathrm{m}^2\,\mathrm{s}^{-1}$ ($l_a = 7.1\,\mu\mathrm{m}$).

### Simulation parameter value estimation

The simulation model presented in Fig. 3A of the main text contains many parameters, which are listed in Table 1. We constrain these parameters by combining several experimental observations.

First, for model simplicity, we chose to have a single lattice spacing between cross-linker binding sites both on the microtubule and on the actin filaments. We take this lattice spacing to be equal to the distance between subunits of the microtubule, which is $\delta = 8\,\text{nm}$ [9]. The actual distance between binding sites on the actin filament is around $5\,\text{nm}$ [10, 11]. However, the actin binding domain of TipAct consists of a tandem of CH domains and TipAct forms a dimer [2], making it likely that TipAct typically fills two neighboring binding sites. Moreover, due to the helical twist of the actin filament [12], it is unclear how many binding sites are available on average. Hence, we can only find an approximate value for the lattice spacing between binding sites on the actin filament. Importantly, our model results do not depend qualitatively on the values of the lattice spacings on the microtubule and the actin filament.

A second parameter that can be estimated relatively easily from experiments is the size of the microtubule tip region. In our model, the tip region is a deterministic step function. In experiments, the EB3-TipAct comet fills roughly a single pixel of the camera, which has a size of $0.2\,\mu\text{m}$. Therefore, we take the length of this tip region $l_t = 0.2\,\mu\text{m}$ as a rough estimate, which sets the number of tip sites to 25 using the previously introduced choice for lattice spacing $\delta$.

Next, the longitudinal friction coefficient of F-actin due to its viscous interactions with the solution can be calculated theoretically for a thin rod [13–15],

$$\zeta_{||} = \frac{2\pi\eta l_a}{\log(l_a/d) + \gamma_{||}}. \tag{10}$$

Here, the correction parameter $\gamma_{||} = -0.114$, as confirmed experimentally for actin filaments [15], and the actin diameter $d = 9 \times 10^{-9}\,\text{m}$ [12]. Furthermore, the viscosity is taken to be that of water, $\eta = 1 \times 10^{-3}\,\text{Pa s}$, and we use a filament length of $l_a = 3\,\mu\text{m}$ here, which falls within the experimental length distribution shown in Fig. 2G. Using the Einstein relation [16] and thermal energy $k_B T = 4.1 \times 10^{-21}\,\text{J}$, this gives us $D_a \approx 1\,\mu\text{m}^2\,\text{s}^{-1}$. For simplicity, we will not vary the value of $D_a$ with the actin length, which is justified by the fact that the cross-linkers limit the motion of the actin filament much more strongly than the viscous drag by the solvent.

Another parameter in our model is $\tau_a$, the time it takes for the actin filament to diffuse far enough away from the microtubule such that it is unlikely to be recaptured. To estimate this time scale, we use the time it takes diffusion of the actin filament to produce a standard deviation of $0.1\,\mu\text{m}$ in the perpendicular position. This is roughly three times the length of the TipAct-EB3 complex [17], and also accounts for the possibility that the actin filament could slightly bend. Using an equation for the friction coefficient $\zeta_\perp$ similar to Eq. 10 [15], we arrive at

$$\tau_a \approx 5 \times 10^{-3}\,\text{s}. \tag{11}$$

As Fig. 3A shows, we have to constrain the rates $r^T_{0,1}$, $r^T_{1,0}$, $r^L_{0,1}$, $r^L_{1,0}$, $r^0_{1,2}$, and $r^0_{2,1}$, and also the spring constant $k$. Since these parameters are difficult to access directly, we estimate them based on a comparison between experimental observations and simulation results. Firstly, we use the observed diffusion constants of actin filaments on the lattice. Secondly, we measure the duration of actin diffusion on the microtubule lattice before unbinding as a function of the actin length. Thirdly, we fit our parameter set to the order of magnitude of the duration of actin transport events. Fourthly, we label TipAct and EB with the same fluorescent tag, and measure the difference in the fluorescence signal between the microtubule lattice and tip regions as a read-out of the binding affinity of the complex. Lastly, we use several specific observations on the (un)binding rates obtained by fluorescence recovery after photobleaching (FRAP) measurements. We explain all these experimental sources in detail below.

We discussed above that the actin diffusion constant is roughly $D_{\text{eff}} = 0.01\,\mu\text{m}^2\,\text{s}^{-1}$ at $l_a = 7\,\mu\text{m}$, as shown in Fig. 5. This diffusion constant is affected by the average number of cross-linkers that bind to the actin filament. If there are many cross-linkers, the diffusion on the lattice slows down. The number of bound linkers is set by all binding rates on the lattice, $r^L_{0,1}$, $r^L_{1,0}$, $r^0_{1,2}$, and $r^0_{2,1}$. The parameter $r^0_{2,1}$ is also important for actin movement in another way, since movement is limited by how fast the cross-linkers can remodel. Hence, a higher $r^0_{2,1}$ will quickly increase the effective diffusion constant of actin on the microtubule lattice by reducing the number of cross-linkers and by increasing the rate of cross-linker remodeling. Additionally, the actin diffusion constant decreases by lowering the spring constant $k$, since this increases the numbers of bound cross-linkers by increasing the accessibility of the actin binding sites.

Another experimental observation that constrains the binding rates is the average duration of actin diffusion on the microtubule lattice. Fig. 5D shows that this duration depends exponentially on the actin length, since it becomes exponentially more likely that at least one cross-linker binds to the actin filament when more binding sites are available. The experimentally observed duration is strongly stochastic, and since we only have 19 events where actin diffuses on the lattice, we can only make an order of magnitude estimate of the average time of being bound for a diffusing actin filament of a certain length. The experiments show that the actin stays on the microtubule lattice for roughly 40 s at $l_a = 4\,\mu\text{m}$, so we fit the order of magnitude of the actin binding time in our simulations to this value. Specifically, we perform simulations in which a 4 µm long actin filament is transported by the microtubule, and then record both the time until the actin falls behind the tip and the time until the actin unbinds from the microtubule lattice. Several parameters influence the duration of lattice diffusion in the simulations. Increasing the average number of cross-linkers on the lattice by changing $r^L_{0,1}$, $r^L_{1,0}$, $r^0_{1,2}$ or $r^0_{2,1}$ makes this binding time larger. Additionally, the actin binding time can be increased by reducing the time scale of cross-linker remodeling, so by increasing both $r^0_{1,2}$ and $r^0_{2,1}$ while keeping the number of cross-linkers fixed. The actin unbinds when no cross-linkers connect the filaments for a duration of $\tau_a$, and rebinding events often rescue the actin filament from unbinding if the cross-linkers remodel quickly.

Then, we use a rough estimate of the average time an actin filament is transported by a growing microtubule tip in cases where microtubule catastrophes and actin unbinding play no role. We used data sets that were limited in number of analysed transport events, which are part of the data presented in Fig. 6AB, Fig. 1E and Fig. 1G, together with Eq. 1 to estimate the average time $T_e \approx 80\,\text{s}$ for an actin filament of $l_a = 4\,\mu\text{m}$ and a microtubule growth velocity of $v_g = 3\,\mu\text{m}\,\text{min}^{-1}$. We hence excluded simulation parameter sets that showed order of magnitude deviations from this value of $T_e$.

All binding parameters have an influence on the transport time $T_e$. Increasing the condensation force leads to longer transport times, and a larger condensation force can be created by increasing the binding affinity on the tip compared to the lattice. This can be seen in Eq. 17, where changing $k$, $r^T_{0,1}$, $r^T_{1,0}$, $r^L_{0,1}$ or $r^L_{1,0}$ changes the condensation force. Increasing the number of cross-linkers by altering $r^0_{1,2}$ and $r^0_{2,1}$ also increases the force, since the stronger interaction between the actin filament and the microtubule amplifies the effect of the stronger affinity for the lattice as set by $r^T_{0,1}$ and $r^T_{1,0}$. However, simply increasing the force by increasing the number of cross-linkers is not generally favorable, since it also increases friction between the moving actin filament and the fixed microtubule.

Another set of experiments shows that TipAct has a significant affinity for the microtubule lattice, which we can directly compare to the affinity for the microtubule tip. Specifically, we observe that the intensity of TipAct and EB3 is roughly $R = 10$ times higher on the tip than on the lattice when no actin is present, as shown in Fig. 3. This ratio is set by the rates $r^T_{0,1}$, $r^T_{1,0}$, $r^L_{0,1}$ and $r^L_{1,0}$,

$$R = \left(\frac{K^t_{0,1}}{1+K^t_{0,1}}\right) / \left(\frac{K^\ell_{0,1}}{1+K^\ell_{0,1}}\right), \tag{12}$$

where the equilibrium constants $K^t_{0,1}$ and $K^\ell_{0,1}$ are defined in Eq. 14. These same parameters also influence the condensation force, as seen in Eq. 17. Hence, the ratio $R$ sets a limit on the condensation force in the limit where $K_{1,2} \to \infty$,

$$F_{f,max} = \frac{k_B T}{\delta} \log[R] \approx 1.2\,\text{pN}. \tag{13}$$

The true force $F_f = 0.10\,\text{pN}$ is more than an order of magnitude lower, since $K_{1,2}$ is far from $\infty$ using our final parameter set. So far, we have only talked about the effects of the ratios $K_{0,1}$, but we do not have data on the specific rates $r^L_{0,1}$ or $r^L_{1,0}$. Hence, we make a choice that makes $r^L_{0,1} < r^T_{0,1}$ and $r^L_{1,0} > r^T_{1,0}$, but the specific choice has little effect as long as the factors $K_{0,1}$ stay the same.

To gain insight into the rate $r^0_{2,1}$ at which TipAct unbinds from the actin while it remains connected to the microtubule, we use *in vitro* TIRF measurements of the TipAct fluorescence signal on single actin filaments, as shown in Fig. 7A. We find that TipAct barely binds to actin filaments when no microtubule is present, and the binding events that do happen are quickly followed by unbinding events. From measurements with an inter-frame duration of $\Delta T = 33\,\text{ms}$, a mean binding time of $\overline{T} = 111\,\text{ms}$ is deduced (Fig. 7A), giving an off-rate of $r_{\text{off}} \approx 9\,\text{s}^{-1}$ from actin without a microtubule present. However, the rate $r^0_{2,1}$ used in the simulations does not simply equal $r_{\text{off}}$. In the simulations, TipAct unbinds from the actin filament but remains bound to a microtubule that is very close. Furthermore, in the experiments shown in Fig. 7A, unbinding means that the cross-linker has not only detached but also that it has diffused away, since it is no longer in the proximity of the actin filament. In the simulations, what is required is only a short moment where the cross-linker is not bound, and we consider a quick rebinding to a neighboring location to be a new binding event. Hence, the binding and unbinding rates in our model will be higher than the $r_{\text{off}}$ observed in full (un)binding experiments.

The (un)binding rates for the TipAct on the microtubule tip also require some interpretation. Since the protein is assumed to bind in a complex with EB3, TipAct and EB3 could in principle unbind individually. In the simulations, we consider only a single cross-linker, ignoring the possibility of the complex dissociating. To estimate the (un)binding rates $r^T_{0,1}$ and $r^T_{1,0}$, kinetic data is available on EB1 [3], and on TipAct and EB3 on microtubule tips as shown in Fig. 7BC. The binding rate of EB1 to the microtubule tip is $0.15\,(\text{nM})^{-1}\text{s}^{-1}$ [3]. With an EB concentration of 100 nM, we would get a binding rate of $15\,\text{s}^{-1}$. However, we assume that the concentration of the complex equals the (limiting) TipAct concentration, which is 25 nM in our experiments.

For the unbinding rate of the complex from the microtubule tip $r^t_{1,0}$, Fig. 7BC shows that we found unbinding rates of $1.8\,\text{s}^{-1}$ for EB3 on microtubule tips, $0.8\,\text{s}^{-1}$ on microtubule tips associated with actin bundles, $3.1\,\text{s}^{-1}$ for TipAct on microtubule tips, and $0.7\,\text{s}^{-1}$ for TipAct cross-linked between microtubules and actin bundles. The reported unbinding rate of EB1 from microtubule tips equals $3.4\,\text{s}^{-1}$ [3]. We need to assume a higher unbinding rate $r^t_{1,0}$, since a rate of $3.4\,\text{s}^{-1}$ makes partial binding very likely and makes it impossible for sites not to be bound. We justify this choice by seeing that this rate is an effective parameter that combines the unbinding of EB3 from the microtubule and of TipAct from EB3. Furthermore, the floppiness of the actin makes the binding of cross-linkers less likely because the actin is occasionally at a distance from the microtubule, but this flexibility is completely ignored in our model. Hence, we overestimate the likelihood of the fully bound state of the cross-linkers, which we can compensate for by increasing $r^t_{1,0}$. An additional reason why $r^t_{1,0} > 3.4\,\text{s}^{-1}$ is that the rate of $3.4\,\text{s}^{-1}$ obtained by FRAP measures the effective unbinding rate after possible rebinding events [3], whereas we consider the short-time unbinding events to be separate events.

The value for the spring constant $k$ of the cross-linkers is also not easily accessible by direct measurements. We choose a value of $2 \times 10^4\,\text{k}_\text{B}\text{T}/\mu\text{m}^2$, which is lower than the range of previously reported spring constants $6 \times 10^4\,\text{k}_\text{B}\text{T}/\mu\text{m}^2$ to $3 \times 10^5\,\text{k}_\text{B}\text{T}/\mu\text{m}^2$ for other cytoskeletal cross-linking proteins [1, 18–20]. This low value accounts for the flexibility of a complex of EB3 dimers and TipAct dimers, and for the floppiness of actin. We chose several values of $k$ going down from $1 \times 10^5\,\text{k}_\text{B}\text{T}/\mu\text{m}^2$ [1] to $1 \times 10^4\,\text{k}_\text{B}\text{T}/\mu\text{m}^2$, then varying the binding rates until we found parameter sets that roughly complied with all considerations listed in this section. The value of $k = 2 \times 10^4\,\text{k}_\text{B}\text{T}/\mu\text{m}^2$ leads to a parameter set where the effective diffusion constant of the cross-linked actin filament is consistent with the experimental measurements. Lowering the spring constant even more leads to frequent events where the actin filament stochastically moves to $x \sim 1\,\mu\text{m}$ in the co-moving reference, making it protrude beyond the front of the growing microtubule tip. An actin filament that sticks so far past the front of the microtubule should be observable in TIRF microscopy. Since we do not observe such events experimentally, we did not lower the spring constant further.

We finally arrive at a set of parameters that accounts for all the experimental observations, listed in Table 1. We find an actin diffusion constant on the microtubule lattice of $D_{\text{eff}} = 0.006\,\mu\text{m}^2\,\text{s}^{-1}$ for an actin length of $7\,\mu\text{m}$, comparable with the experimental values of $0.014(3)\,\mu\text{m}^2\,\text{s}^{-1}$ ($l_a = 6.6\,\mu\text{m}$) and $\overline{D} = 0.009(3)\,\mu\text{m}^2\,\text{s}^{-1}$ ($l_a = 7.1\,\mu\text{m}$). The average transport duration is 90 s in simulations for a microtubule growth velocity of $3\,\mu\text{m}\,\text{min}^{-1}$ and an actin length of $4\,\mu\text{m}$, consistent with the experimental value of 80 s. With the final parameter set, an actin filament of $4\,\mu\text{m}$ unbinds from the microtubule roughly 44 s after it loses its interaction with the microtubule tip in the simulations, close to the experimental value of 40 s observed in the actin diffusion experiments. Finally, our parameter set leads to a factor $R = 26.1$ from Eq. 12, which corresponds to an error in the affinity of less than $1\,\text{k}_\text{B}\text{T}$ compared to the factor $R = 10$ measured experimentally.

### Analytical expression for condensation force

The magnitude of the condensation force that drives actin transport by microtubule tips depends on the strength of the interaction between the cross-linkers and the filaments and on the density of binding sites in the overlap between the actin filament and the microtubule tip region. Here we derive an analytical expression for this force.

Because the model does not allow cross-linkers that are only bound to the actin filament, we focus on the binding sites on the microtubule and classify the sites according to their cross-linker binding state, as shown in Fig. 8. The state is denoted by 0 if the site is free, 1 if the microtubule site is occupied by a cross-linker that is dangling and not connected to the actin, and 2 if the site is occupied by a cross-linker that is also bound to the actin. We use $r_{i,j}$ for the transition rate from state $i$ to state $j$, as shown in Fig. 3A of the main text. We define the equilibrium constants $K_{i,j}$ between the states through the local detailed balance relation,

$$\begin{aligned} K^{\alpha}_{0,1} &= \exp(-\left(\mathcal{F}^{\alpha}_{1} - \mathcal{F}_0\right)/k_BT) = \frac{r^{\alpha}_{0,1}}{r^{\alpha}_{1,0}}, \\ K_{1,2} &= \exp(-\left(\mathcal{F}^{\alpha}_{2} - \mathcal{F}^{\alpha}_{1}\right)/k_BT) = \frac{r^{0}_{1,2}}{r^{0}_{2,1}}\sqrt{\frac{2\pi k_BT}{k\delta^2}}, \end{aligned} \tag{14}$$

where $\alpha \in \{t, \ell\}$ represents the microtubule tip or lattice region, respectively. The equilibrium constants are ratios of partition functions, and are thus related to differences between the free energies $\mathcal{F}_i$ of different binding states $i$. The process to create a cross-linker in the dangling state 1 depends on the microtubule region $\alpha$ that the cross-linker is binding to, because cross-linkers bind faster to and unbind slower from the microtubule tip than from the lattice. However, we assume that the rate to bind to the actin filament is the same for all dangling cross-linkers, independent of the microtubule region they are bound to. Hence, the equilibrium constant $K_{1,2}$ does not depend on $\alpha$. The rate $r^{\alpha}_{0,1}$, and hence $K^{\alpha}_{0,1}$, is proportional to the TipAct concentration in the solution, but we keep this dependence implicit since we do not vary the concentration in the experiments. Since the cross-linkers act as harmonic springs, a cross-linker bound to a specific binding site on the microtubule can bind to multiple binding sites on the actin filament. We group all these possible binding states into a single state 2. Therefore, the expression for $K_{1,2}$ is a sum over all possible cross-linker binding positions on the actin filament, and we approximate the sum by assuming that there are no interactions between the cross-linkers and that the actin filament is infinitely long,

$$K_{1,2} \approx \sum_{i=-\infty}^{\infty} \frac{r^{0}_{1,2}}{r^{0}_{2,1}} \exp\left(-\frac{k\,(i\delta)^2}{2k_BT}\right) \approx \int_{-\infty}^{\infty} \frac{1}{\delta}\frac{r^{0}_{1,2}}{r^{0}_{2,1}} \exp\left(-\frac{kx^2}{2k_BT}\right) \mathrm{d}x = \frac{r^{0}_{1,2}}{r^{0}_{2,1}}\sqrt{\frac{2\pi k_BT}{k\delta^2}} \tag{15}$$

Here, we make use of our model assumption that the actin filament has lattice spacing $\delta$.

To calculate the condensation force, we consider the case where an actin filament overlaps with both the microtubule lattice and tip regions, such that the front of the actin filament is behind the front of the microtubule, as shown in Fig. 8. We assume that there are $n_\ell$ microtubule lattice sites and $n_t$ microtubule tip sites overlapping with the actin filament, such that the actin filament contains $\ell_a = n_\ell + n_t$ sites. Furthermore, the microtubule has a total of $\ell_\ell$ lattice sites and $\ell_t$ tip sites. For the calculation of the force, we also assume that the cross-linker binding states of different microtubule binding sites are independent, such that the partition function describing the binding state of the full system can be expressed as the product of the local partition functions for each site on the microtubule. The microtubule sites can always be in states 0 and 1, but state 2 is only available when the actin filament overlaps with that site. The local partition function for each microtubule binding site is a sum over the possible binding states of the Boltzmann factors. These factors contain the free energies $\mathcal{F}_i$, and we make the choice to set the free energy $\mathcal{F}_0 \equiv 0$, giving a Boltzmann factor of 1 for state 0. Using Eq. 14, we see that the Boltzmann factors for binding states 1 and 2 are simply products of the equilibrium constants. Hence, the full partition sum for the system shown in Fig. 8 is

$$\mathcal{Z}(n_\ell, n_t) = \left[1 + K^\ell_{0,1}\right]^{\ell_\ell - n_\ell} \left[1 + K^\ell_{0,1} + K^\ell_{0,1} K_{1,2}\right]^{n_\ell} \left[1 + K^t_{0,1} + K^t_{0,1} K_{1,2}\right]^{n_t} \left[1 + K^t_{0,1}\right]^{\ell_t - n_t} . \quad (16)$$

The four factors correspond to the $(\ell_\ell - n_\ell)$ lattice sites outside of the actin overlap, the $n_\ell$ lattice sites within the actin overlap, the $n_t$ tip sites within the actin overlap, and the $(\ell_t - n_t)$ tip sites outside of the actin overlap, as shown in Fig. 8A. The condensation force can be calculated as a positional derivative of the free energy $\mathcal{F}(n_\ell, n_t) = -k_B T \log[\mathcal{Z}(n_\ell, n_t)]$, given by Eq. 16. The actin filament position $x$ is defined in Fig. 3A of the main text as the difference between the front of the actin filament and the location on the microtubule where the lattice region turns into the tip region, and Fig. 8A shows that this region contains $n_t$ sites that are spaced $\delta$ apart. Hence, $n_t = x/\delta$, and since the number of sites on the actin filament $\ell_a$ is constant, $n_\ell = \ell_a - n_t = \ell_a - x/\delta$. The forward pointing condensation force can thus be calculated as

$$F_f = -\frac{\mathrm{d}\mathcal{F}(\ell_a - x/\delta, x/\delta)}{\mathrm{d}x} = \frac{k_B T}{\delta} \log\left[\frac{1 + K^\ell_{0,1}}{1 + K^\ell_{0,1} + K^\ell_{0,1} K_{1,2}} \frac{1 + K^t_{0,1} + K^t_{0,1} K_{1,2}}{1 + K^t_{0,1}}\right] . \quad (17)$$

We see that the condensation force does not depend on the position of the actin filament $x$ as long as there is a partial overlap with the microtubule tip region. The situation changes when $x > l_t = \ell_t \delta$, such that the actin fully covers the microtubule tip region and sticks out in front of the microtubule. Then, the *forward* pointing condensation force disappears because it is no longer possible for the actin filament to gain binding free energy by increasing the overlap with the microtubule tip. Further forward motion of the actin filament now leads to a loss of the number of binding sites on the microtubule lattice, which comes with a free energy cost. In this situation, while the number of tip sites overlapping with the actin filament $n_t = \ell_t$ remains constant, $n_\ell$ still depends on $x$ as before, and Eq. 16 shows that there will be a *backward* pointing condensation force due to the loss of overlap with the microtubule lattice,

$$F_b = -\frac{\mathrm{d}\mathcal{F}(\ell_a - x/\delta, \ell_t)}{\mathrm{d}x} = \frac{k_B T}{\delta} \log\left[\frac{1 + K^\ell_{0,1}}{1 + K^\ell_{0,1} + K^\ell_{0,1} K_{1,2}}\right] . \quad (18)$$

$F_b$ is negative because all equilibrium constants $K$ are positive. Since the binding affinity of cross-linkers for the microtubule lattice is much lower than for the microtubule tip, $K^\ell_{0,1} \ll K^t_{0,1}$, we have that $|F_b| \ll |F_f|$. Using the parameter values of Table 1, we find the values $F_f = 0.10\,\mathrm{pN}$ and $F_b = -4.6\,\mathrm{fN}$, showing that the backward force is typically irrelevant. The condensation force is plotted as a function of the actin position in Fig. 8B. The backward force is only able to move the actin filament when the microtubule is depolymerizing, an effect that we confirmed experimentally (see Fig. 2D). Depolymerizing microtubules have no tip region where EB3 binds strongly, leading to a low number of cross-linkers and thus to a low cross-linker induced friction force. A force in the femtonewton range can easily overcome the viscous friction between the actin filament and the solution, since $F_f$ applied to an actin filament with a diffusion constant of $1\,\mu\mathrm{m}^2\,\mathrm{s}^{-1}$ would lead to a drift velocity of $-1.1\,\mu\mathrm{m}\,\mathrm{s}^{-1}$.

### Condensation force in the optical tweezer experiments

In the optical tweezer experiments, the bead plays the role of the actin filament that is transported by the growing microtubule tip region in the actin transport experiments. Below, we theoretically estimate the condensation force associated with the geometry of the optical tweezer experiment. Because TipAct (in complex with EB3) has a much higher affinity for microtubules than for actin filaments, we assume in the simulations of actin transport that TipAct never binds to the actin filament directly, but that it first binds to the microtubule before it can bind to the actin filament. By contrast, TipAct is irreversibly bound to the bead in the optical tweezer experiments, and there is no TipAct in solution. Hence, the only possible binding path is from the dangling state on the bead to a fully connected state between the bead and the microtubule. The two distinct regions of the microtubule then lead to two equilibrium constants describing the affinity for the microtubule, $K_t$ and $K_\ell$. An additional key parameter determining the force is the density of bound cross-linkers on the bead $\rho$.

Calculating the free energies relative to the dangling state, such that this state has the Boltzmann factor 1, we derive the force on the bead using a similar method as used to derive Eq. 17,

$$F_{f,\text{bead}} = k_B T \rho \log\left[\frac{1+K_t}{1+K_\ell}\right]. \tag{19}$$

We do not have experimental data to estimate these parameters accurately, but we can make an order of magnitude estimate. First, we estimate the density of cross-linkers on the bead $\rho$. This density represents the number of cross-linkers that is conformationally capable to bind per micrometer of microtubule length. We make the order of magnitude estimate that $\rho$ equals 10% of a fully covered microtubule, $\rho \approx 0.1/\delta$.

To find the equilibrium constants $K_t$ and $K_\ell$, we recognize that TipAct binds much more strongly to microtubules than to single actin filaments, since single microtubule tips show a strong fluorescent TipAct signal while single actin filaments do not (Fig. 1B, Fig. 1C). Measuring dwell times (Fig. 7A) and performing fluorescent recovery after photobleaching (FRAP) measurements (Fig. 7B), we report dissociation constants of $K_D^t = 67\,\text{nM}$ for the binding between TipAct and microtubule tip regions, and $K_D^a = 5.2\,\mu\text{M}$ for the binding between TipAct and actin [17]. We note that $K_D^a$ was probably underestimated in the experiments, because TipAct forms dimers that have two actin binding sites, which likely caused actin filaments to bundle and led to a higher measured affinity than should be expected for the binding of TipAct to single actin filaments. To find a relation for the parameters $K_t$ and $K_\ell$, we use that the equilibrium constant $K_t$ describes the transition to bind from the bead (representing the actin filament) to the microtubule, and $K_{1,2}$ represents the transition to bind from the microtubule to the actin filament. Based on the experimental values, we have $K_t/K_{1,2} \approx K_D^a/K_D^t = 78$, giving us the order of magnitude $K_t = 100K_{1,2}$ in the simulations. This is likely an underestimation due to the described bias in the value of $K_D^a$.

Finally, to find $K_\ell$, we use that the difference in affinity between the microtubule tip and lattice regions should be the same for the optical tweezer experiments and the actin transport experiments, $K_t/K_\ell = K_{0,1}^t/K_{0,1}^\ell$. Using Table 1, we find the force

$$F_{\text{f,bead}} \approx 0.17\,\text{pN}. \tag{20}$$

This is the force on the bead when the bead is behind the front of the microtubule and still overlapping with the microtubule tip region. However, when the bead and the microtubule first come into contact, the bead is in front of the microtubule. Then, the condensation force caused by the increasing overlap with the microtubule tip region equals

$$F_{b,\text{bead}} = -k_B T \rho \log[1+K_t] \approx -0.21\,\text{pN}, \tag{21}$$

using the same parameters as before. As shown in Fig. 9, we experimentally find a forward force of $F_{\text{f,bead}}0.12\,\text{pN}$ and a backward force of $F_{\text{b,bead}} \approx -0.7\,\text{pN}$. Hence, we correctly estimate the order of magnitude for these forces, but the ratio between these forces is not well predicted. This may be due to an overestimation of the ratio $K_t/K_\ell$, which is in turn due to an overestimation of $K^t_{0,1}/K^\ell_{0,1}$.

### Probability of actin unbinding

In our simulations, we consider the actin filament to be bound to the microtubule as long as there is at least a single cross-linker binding the two filaments together. Here, we obtain an approximate expression for this binding probability using similar expressions as in the derivation of the condensation forces. The probability of having no cross-linkers connecting an actin filament that overlaps with the microtubule tip region over a length $y_t$ and overlaps with the microtubule lattice region over a length $y_\ell$, $p_0(y_t, y_\ell)$ defined in Eq. 5 of the main text, equals

$$p_0(y_t, y_\ell) = \left[\frac{1 + K^t_{0,1}}{1 + K^t_{0,1} + K^t_{0,1}K_{1,2}}\right]^{y_t/\delta} \left[\frac{1 + K^\ell_{0,1}}{1 + K^\ell_{0,1} + K^\ell_{0,1}K_{1,2}}\right]^{y_\ell/\delta} = \mathrm{e}^{-y_t/\lambda_t}\mathrm{e}^{-y_\ell/\lambda_\ell}. \tag{22}$$

Here, the discrete numbers of binding sites that overlap with the microtubule tip region and lattice region are approximated as $n_t = y_t/\delta$ and $n_\ell = y_\ell/\delta$, respectively. We recognize that the probability decreases exponentially with both overlap lengths, which allows us to define the length scales $\lambda_t$ and $\lambda_\ell$,

$$\lambda_\alpha = \frac{\delta}{\log\left[1 + K^\alpha_{0,1} + K^\alpha_{0,1}K_{1,2}\right] - \log\left[1 + K^\alpha_{0,1}\right]}, \tag{23}$$

with $\alpha \in \{t, \ell\}$ representing the microtubule region again. Using the parameters listed in Table 1, we calculate $\lambda_t = 0.038\,\mu\mathrm{m}$ and $\lambda_\ell = 0.89\,\mu\mathrm{m}$. Since there is a higher density of bound cross-linkers on the microtubule tip than on the lattice, as shown in Fig. 3, the probability to have no cross-linkers decays much more rapidly with an increase in the overlap length in case of the tip region as opposed to the lattice region. This explains why the length scale $\lambda_t$ is much shorter than $\lambda_\ell$.

### Kramers theory

In the main text, we identify that an actin filament that is being transported by a growing microtubule needs to cross a free-energy barrier to fall behind the microtubule tip region. Given that we know the free-energy profile and the diffusion constant, we can calculate the rate of these transitions using Kramers theory.

As shown in Fig. 3C of the main text, $x$ is the position of the front of the actin filament relative to the point on the microtubule where the lattice region ends and the tip region begins. Since the latter point moves with the growing microtubule tip, $x$ constitutes a co-moving frame of reference. In this frame of reference, we call $D(x)$ the effective diffusion constant of the actin filament. We obtain $D(x)$ by calculating how the overlaps between the actin filament and the microtubule lattice region $y_\ell$ and between the actin filament and the microtubule tip region $y_t$ change with $x$. As shown in Fig. 3C of the main text,

$$\begin{aligned} y_t(x) &= x\mathbb{1}(0 \le x < l_t) + l_t\mathbb{1}(x \ge l_t)\,, \\ y_\ell(x) &= l_a - x\mathbb{1}(x \ge 0)\,, \end{aligned} \tag{24}$$

where we assume that $l_a \gg l_t, x$. Then, we enter these expressions into Eqs. 2&3 of the main text to find $D(x) = D(y_t(x)\,, y_\ell(x))$, which is given in Eq. 1 of the main text.

Next, we call $F_{\text{eff}}(x)$ the forward pointing force on the actin filament, given by Eq. 4 of the main text and containing a contribution from the friction. We integrate this effective force to find the generalized free energy

$$\mathcal{F}(x) = -\int_0^x F_{\text{eff}}(x')\,\mathrm{d}x'. \tag{25}$$

The function $\mathcal{F}(x)$ behaves exactly like an equilibrium free energy, even though it describes the non-equilibrium process of microtubule growth and actin transport. This is the result of the co-moving frame, in which the free-energy profile loses its time dependence.

Combining the expressions for the diffusion constant and the generalized free energy, and assuming the Einstein relation [16], we find a Fokker-Planck equation for $f_a(x,t)$, the probability density of the actin position $x$,

$$\partial_t f_a(x,t) = \partial_x \left[ D(x)\, \mathrm{e}^{-\mathcal{F}(x)/k_B T} \partial_x \left( \mathrm{e}^{\mathcal{F}(x)/k_B T} f_a(x,t) \right) \right]. \tag{26}$$

As shown in Eqs. 1–4 of the main text, the free energy is determined by the parameters $D_a$, $\lambda_t$, $\lambda_\ell$, $\zeta_t$, $\zeta_\ell$, $F_f$, and $F_b$. $D_a$ is a parameter in the simulations as well, and we have analytical approximations to the length scales $\lambda_t$ and $\lambda_\ell$ in Eq. 23, and for the condensation forward and backward forces $F_f$ and $F_b$ in Eq. 17 and Eq. 17, respectively. The only fitting parameters are the two proportionality factors $\zeta_t$ and $\zeta_\ell$ that determine how the friction coefficient scales with the actin length on the microtubule tip and lattice. Direct measurements of the diffusion constant in simulations, shown in Fig. 3D of the main text, give the values $\zeta_t = 810\,\mathrm{k_B Ts/\mu m^3}$ and $\zeta_\ell = 30.2\,\mathrm{k_B Ts/\mu m^3}$. We emphasize that this set of simulations is independent of the set of simulations in which we estimate the free-energy profile and transport times, which means that we can make independent theoretical predictions on the generalized free-energy profile and on the functional behavior of the actin transport time. We obtain the rate of actin falling behind the tip using Kramers theory [21] based on Eq. 26, and the resulting rate is given in Eq. 6 of the main text.

### Rate of actin unbinding

One of the processes by which actin transport ends is by the direct unbinding of the actin filament from the microtubule when it is still in contact with the microtubule tip region. Here, we develop a theoretical explanation of the unbinding rate observed in the simulations. If the actin filament is at a position $x < 0$, transport will most likely rapidly end with the filament falling further behind the tip, which we treat as an process independent from unbinding. Hence, we consider the scenario $x > 0$ for the unbinding rate, such that the actin filament overlaps both with the microtubule lattice region and with the tip region.

The actin filament can only unbind when there are no cross-linkers binding the actin filament. First, the actin filament needs to loose all connections with the microtubule lattice region. Second, the cross-linkers bind very strongly to the microtubule tip region, so the actin can only unbind when the overlap between the actin filament and the tip region almost vanishes, $x \approx 0$. Then, we assume that the rate of unbinding is limited by the probability that both these events happen,

$$r_u \propto \mathbb{P}[\text{no connections lattice} \wedge x \approx 0] = \mathbb{P}[\text{no connections lattice}]\, \mathbb{P}[x \approx 0 \mid \text{no connections lattice}]. \tag{27}$$

Here, we use the definition of conditional probability to rewrite the proportionality into two factors, and we will discuss each factor separately.

First, we calculate the probability that no cross-linkers are bound between the actin filament and the microtubule lattice using Eq. 22. This probability limits the rate at which the actin filament can unbind in Eq. 27, so the unbinding rate decreases exponentially with the actin filament length,

$$r_u \propto \mathrm{e}^{-l_a/\lambda_\ell}. \tag{28}$$

We only focus on how the unbinding rate is shaped by the actin length $l_a$ and the microtubule growth velocity $v_g$, since these are the variables we use to compare the transport time between simulations and experiments. We absorb all other effects into the proportionality constant. We use the length scale $\lambda_\ell$ here, since elongating an actin filament at fixed position $x$ only increases the overlap with the microtubule lattice region, not with the tip region.

Then, we require the probability that $x \approx 0$ given that the actin filament has lost its connections with the microtubule lattice region. This probability follows from the free-energy profile $\mathcal{F}(x)$ shown in Fig. 3E of the main text. Specifically, the peak of the barrier is always located at $x = 0$, while the valley is at $x = l_t$, where $l_t$ is the length of the microtubule tip region. Hence, the probability to find the actin filament close to the peak is given by the free-energy barrier height $\Delta\mathcal{F}^\ddagger = \mathcal{F}(0) - \mathcal{F}(l_t)$,

$$r_u \propto \mathrm{e}^{-\Delta\mathcal{F}^\ddagger/k_BT}. \tag{29}$$

Eq. 25 shows that this free-energy barrier requires integrating the effective force defined in Eq. 4 of the main text. The only term that depends on $v_g$ or $l_a$ is the friction term,

$$F_{\text{eff}}(x) = -\zeta(x)\,v_g + \dots. \tag{30}$$

Given that there are no cross-linkers connecting the actin filament to the microtubule lattice, only the overlap with the microtubule tip provides a significant friction coefficient $\zeta$ for the actin filament,

$$\zeta(x) \approx \zeta_t x. \tag{31}$$

Now, integrating Eq. 30 gives us

$$\Delta\mathcal{F}^\ddagger = \frac{1}{2}\zeta_t l_t^2 v_g + \dots. \tag{32}$$

By combining Eq. 28, Eq. 29, and Eq. 32, we find the actin unbinding rate

$$r_u(v_g, l_a) = r_u^0 \mathrm{e}^{-l_a/\lambda_\ell}\mathrm{e}^{v_g/\gamma}, \tag{33}$$

where the velocity $\gamma$, which sets the scale for the micortubule growth velocity $v_g$, is defined as

$$\gamma = \frac{2k_BT}{\zeta_t l_t^2}. \tag{34}$$

### Supplementary Tables

| Parameter | Value | Sources |
|---|---|---|
| Lattice spacing binding sites $\delta$ | 0.008 µm | literature [2], [9–12]; |
| Microtubule tip size $l_t$ | 0.2 µm | imaging; |
| Binding rate to microtubule tip $r^T_{0,1}$ | 3 s$^{-1}$ | literature [3, 17], transport; |
| Unbinding rate from microtubule tip $r^T_{1,0}$ | 4 s$^{-1}$ | FRAP, transport, literature [3, 17]; |
| Binding rate to microtubule lattice $r^L_{0,1}$ | 1 s$^{-1}$ | fluorescence, diffusion, transport, unbinding; |
| Unbinding rate from microtubule lattice $r^L_{1,0}$ | 60 s$^{-1}$ | fluorescence, diffus., transport, unbind.; |
| Basic binding rate to actin filament $r^0_{1,2}$ | 75 s$^{-1}$ | diffusion, transport, unbinding; |
| Basic unbinding rate from actin filament $r^0_{2,1}$ | 300 s$^{-1}$ | diffusion, transport, unbinding; |
| Effective spring constant $k$ | $2 \times 10^4 k_B T$ µm$^{-2}$ | literature [1], [18–20], diffusion, transport, unbind.; |
| Diffusion constant bare actin filament $D_a$ | 1 µm$^2$s$^{-1}$ | literature [12], [13–15], viscosity; |
| Actin unbinding time $\tau_a$ | $5 \times 10^{-3}$ s | viscosity; |

**Table 1: Model parameters, their values used in the simulations, and the sources used for determining the parameter values.** As explained in the supplemental text, we use a combination of experimental observations to self-consistently choose the model parameters, and most of these observables are influenced by multiple parameters. Hence, we manually varied the parameters and searched for a parameter set that is consistent with all experimental observations. The sources that influenced each parameter value are previously published data (literature), direct microscopy observations (imaging), the typical duration of actin transport events without the effects of actin unbinding or microtubule catastrophe (transport), fluorescence recovery after photobleaching experiments determining effective unbinding rates (FRAP), experimental observations of the actin diffusion constant (diffusion) or binding duration (unbinding) on the microtubule lattice, and calculations of the viscous drag of the solution on an actin filament (viscosity, Eq. S11).

| | MT growth velocity and transport time | | Actin filament length and transport time | |
|---|---|---|---|---|
| | Correlation | Permutation | Correlation | Permutation |
| Experiments (n=265) | −0.16 | $0.01 \pm 0.06$ | 0.17 | $0.01 \pm 0.06$ |
| Experiments selective ends (n=103) | −0.25 | $0.00 \pm 0.09$ | 0.33 | $0.00 \pm 0.09$ |
| Simulations (n=60000) | −0.252 | $0.000 \pm 0.004$ | 0.113 | $0.000 \pm 0.004$ |

**Table 2: Spearman correlation coefficients show that the transport time observed experimentally decreases with the microtubule growth velocity while it increases with the actin length.** By making 100 random permutations of the transport times linked to the growth velocities or the actin lengths, we find the range of Spearman coefficients that can be expected for an uncorrelated data set (mean ± SD), showing that the measured correlation coefficients are significantly outside of this range. By selecting only those events that end by actin falling behind the microtubule tip region ('selective ends' in the table), we find similar correlation coefficients. Finally, we perform the same analysis on the two simulation data sets shown in Figs. 4A,B. Each data point in those figures is the result of simulating 2000 events, giving 66 000 events for the set varying the growth velocity and 60 000 events for the set varying the actin length. The correlation coefficient between the transport time and the growth velocity obtained in the simulations is comparable to the correlation coefficient found in experiments, while the correlation coefficient between the transport time and the actin length agrees in the sign, but is lower in magnitude for the simulations than for the experiments. The reason for this discrepancy is that the simulation data extends to larger actin lengths where the transport time decreases again.

### Supplementary Figures

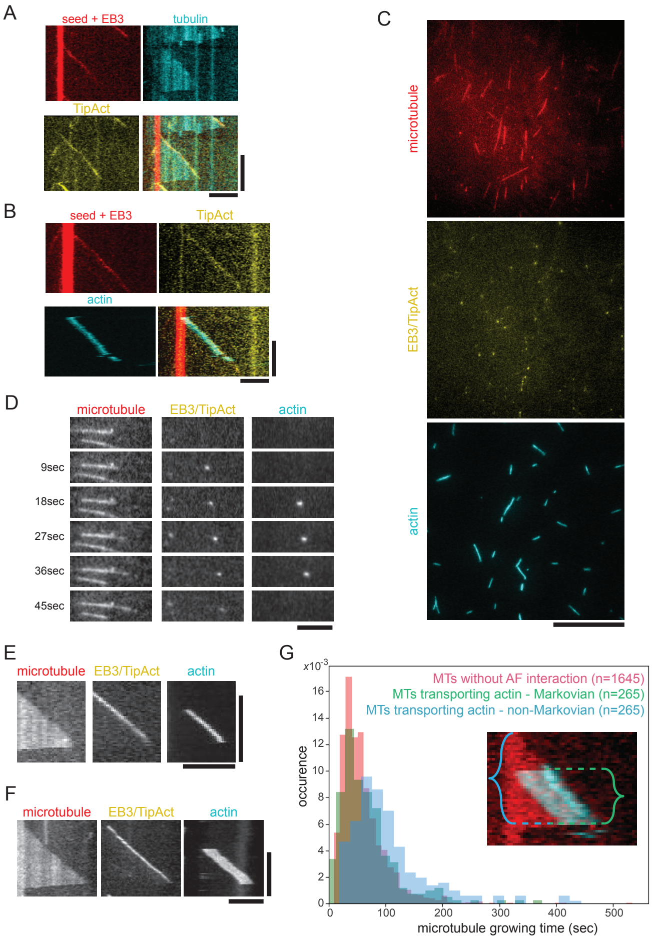


**Figure 1: Supplemental for main Figure 1.** (A) TipAct in the absence of actin filaments tracks the growing microtubule plus tip. Images show microtubule seeds and EB3 both in red (although EB3 does not colocalize to the MT seed, as visible in C), tubulin in cyan, TipAct in yellow, and the merged image. TipAct cannot track microtubule plus-ends independently of EB3 (also shown in [2]. (B) TipAct is present at the growing microtubule plus tip during actin transport events. Images show microtubule seeds and EB3 both in red (although EB3 does not colocalize to the MT seed, as visible in C), TipAct in yellow, actin filaments in cyan, and the merged image. Note that in this experiment, tubulin is not labeled. (C) Separate channels for typical field of view shown in Fig. 1B. A cropped region of 54 by 54 µm was used for analysis. (D-F) Individual channels of transport events shown in main text Fig 1. (D) Time series of the growing plus ends of a microtubule that recruits and transports an actin filament via EB3/TipAct complexes, as is shown in Fig. 1C. (E) Kymograph (a space-time plot) of actin transport event, as is also shown in Fig. 1D (top). (F) Kymograph (a space-time plot) of actin transport event, as is also shown in Fig. 1D (bottom). (G) Normalized distribution of microtubule growing times for growth events where the microtubule does not interact with actin filaments (red) and for growth events where microtubules transport an actin filament (blue). For the actin-transporting microtubule growth events, we also show the growing time starting at the binding of the actin filament (green, a Markovian process), as is indicated by the accolades in the inset too. Average catastrophe rates of 1.00±0.03, 0.89±0.05 and 0.57±0.03 events/min for non-interacting MTs, actin-transporting MTs measured upon actin binding, and complete growth events of actin-transporting MTs, respectively. Number of measured growth events indicated in figure. Protein labels: TipAct is always GFP-labelled. EB3 is mCherry-labelled (in A-B) or GFP-labelled (C-F). Scale bars: 5 µm (horizontal) and 2 min (vertical) in A,B,E and F; 20 µm in C; and 5 µm in D.

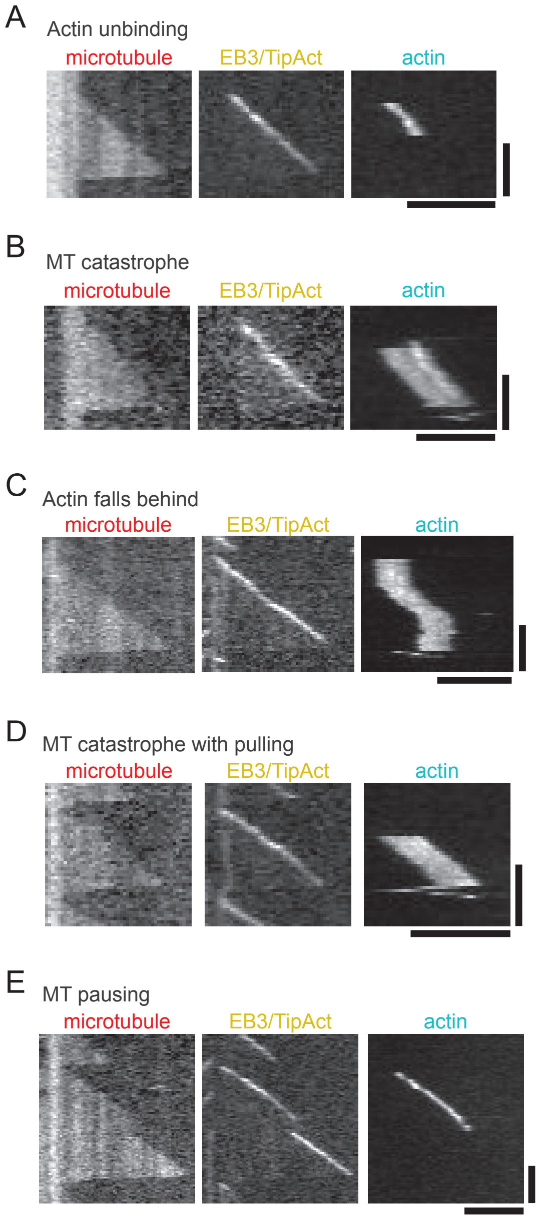


**Figure 2: Individual channels of transport events shown in main Figure 2.** (A) Kymograph of a typical transport event that ends by the unbinding of the actin filament, as is also shown in Fig. 2A. This is the same transport event that is also depicted as a time series in Fig. 1C and Fig. 1D. (B) Kymograph of a typical transport event that ends upon a microtubule catastrophe, as is also shown in Fig. 2B. (C) Kymograph of a typical transport event that ends by loss of contact of the actin filament with the tip resulting in the actin filament falling behind and lingering on the MT lattice, as is also shown in Fig. 2C. (D) Kymograph of a typical transport event that ends upon a microtubule catastrophe where the actin filaments is pulled along by the shrinking microtubule, as is also shown in Fig. 2D. (E) Kymograph of a typical transport event that ends upon a microtubule pausing, as is also shown in Fig. 2E. This is the only example of transport ended by microtubule pausing. In all panels, both EB3 and TipAct are GFP-labelled. Scale bars: 5 µm (horizontal) and 60 s (vertical).

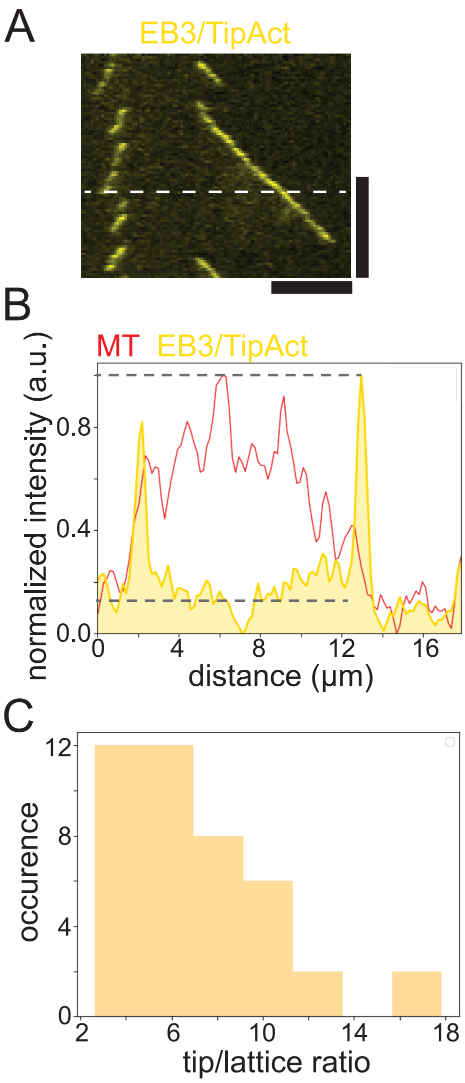


**Figure 3: Ratio between the signal of EB3 and TipAct on the microtubule tip and on the lattice.** (A) Example kymograph showing the (GFP) signal of both EB3 and TipAct, from which tip/lattice-ratios can be calculated. The dashed line was used to produce an intensity profile. (B) Intensity profile showing the lattice (red) and plus end intensity of EB3 and TipAct (yellow). Dashed lines on the profile, showing the maximum value and the mean value along the microtubule lattice for GFP, were used to estimate the 1:10 lattice:tip ratio of the tiptracking-complex. (C) Distribution of tip/lattice ratios for 42 analysed profiles of EB3/TipAct intensity while transporting an actin filament, with 6.9 and 5.9 as the mean and median values of this distribution, respectively. Protein labels: both EB3 and TipAct are GFP-labelled. Scale bars: 5 μm (horizontal) and 2 min (vertical)

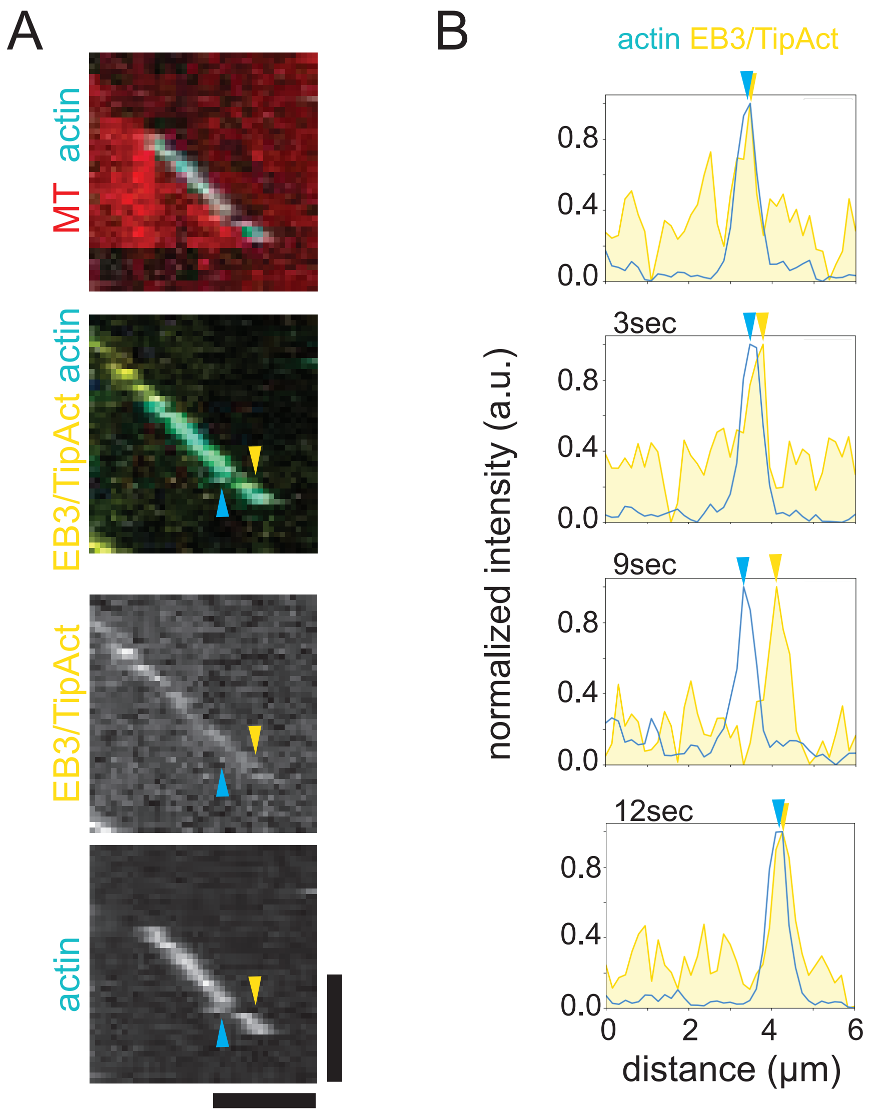


**Figure 4: Catching-up of actin filament to tip-tracking-complex.** (A) Kymograph showing a growing microtubule (red), the tip-tracking-complex consisting of EB3 and TipAct (both yellow), and a transported actin filament (cyan). At one point, the actin filament (cyan arrow) falls behind the tip (yellow arrow), but remains at the microtubule lattice and quickly catches up with the tip-tracking-complex again to continue transport. (B) Line profiles along the kymograph in *A*, showing an actin filament catching up with the tip-tracking-complex. Arrows indicate the locations of the actin filament (cyan) and of the tip-tracking-complex (yellow). Protein labels: both EB3 and TipAct are GFP-labelled. Scale bars: 3 µm (horizontal) and 60 s (vertical).

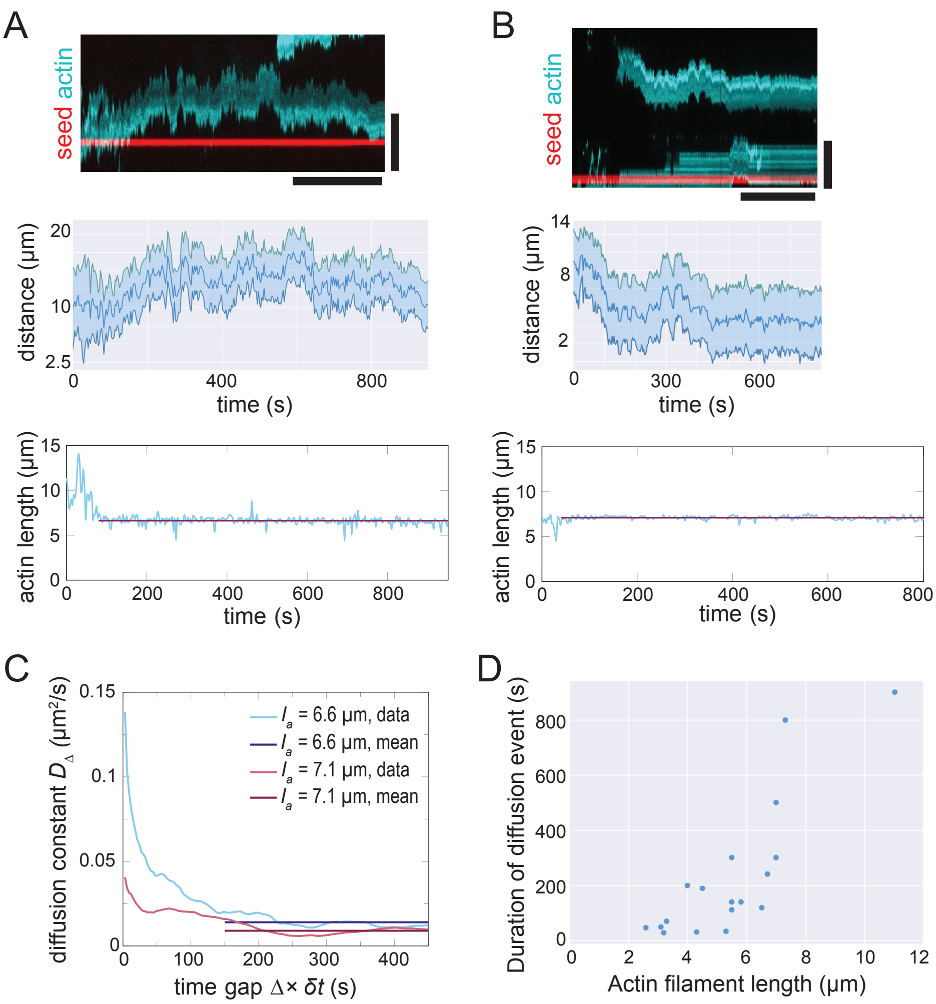


**Figure 5: Diffusion of actin filaments along the microtubule lattice.** (A) Diffusion of 7.1 µm long actin filament along the microtubule lattice in a kymograph (top), corresponding time trace of the filament front and rear end positions (middle), and the filament length fluctuations used to determine the actin filament length (bottom). (B) Another example, showing diffusion of a 6.6 µm long actin filament. (C) Diffusion constant estimates given by Eq. 9. For small time scales, the observed actin position fluctuations are influenced by external noise and length fluctuations, so we can only extract the diffusion constant at large time scales. We take the sample mean of the estimator over the window between 150 s and 450 s, yielding $\overline{D} = 0.014(3)\,\mu m^2\,s^{-1}$ ($l_a = 6.6\,\mu m$) and $\overline{D} = 0.009(3)\,\mu m^2\,s^{-1}$ ($l_a = 7.1\,\mu m$). (D) The typical duration of diffusion events increases with the actin filament length, since the rate at which the actin filament unbinds from the microtubule lattice region decreases with the number of cross-linkers that connect the two filaments. Scale bars in panels A,B: 5 min (horizontal), 10 µm (vertical).

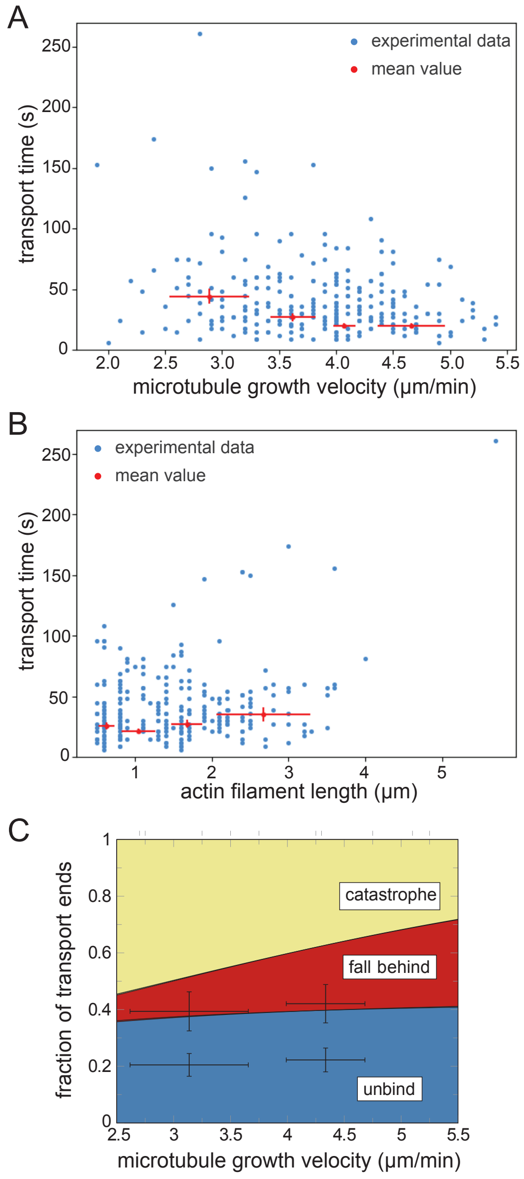


**Figure 6: Unbinned and binned experimental data for actin transport.** (A) The transport time versus the microtubule growth velocity and (B) the transport time versus the actin filament length. This data is complementary to the data shown in main text Fig. 4A,B. Single data points are shown in blue; binned experimental means (when distributing data in 4 bins of equal size) are shown in red and are also depicted in main text Fig. 4A,B. Horizontal error bars represent the standard deviation of the growth velocities or the actin filament length within the bin, and vertical error bars represent the standard deviation of the mean transport time. (C) The fractions of the categories of transport ends as a function of the microtubule growth velocity. The fractions were calculated from experimental values of the catastrophe rate (constant $r_c$), falling behind rate (Eq. 6 of the main text) and unbinding rate (Eq. 5 of the main text). Experimental data was binned into two bins of equal size, shown as points with errors. Transport events ending by microtubule catastrophes include both events where actin is pulled back with the depolymerizing microtubule and events where the actin unbinds upon a catastrophe. The horizontal error bars show the weighted standard deviation of the microtubule growth velocity and of the actin filament length, and the vertical error bars show the standard error of the estimated mean ratio. Since we have to estimate two data points per bin with sufficient statistics, we only divided the growth velocities and actin lengths into two bins.

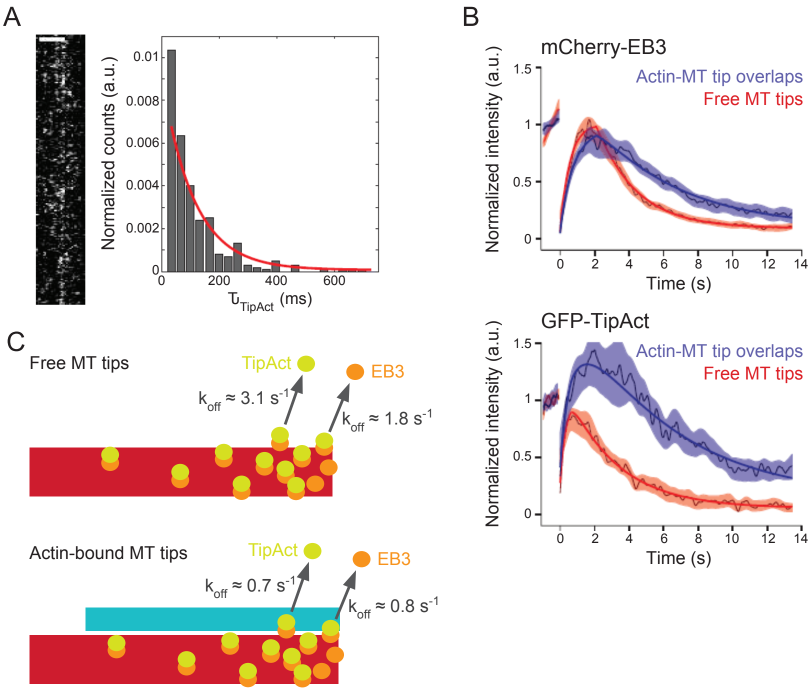


**Figure 7: Binding dynamics of the tip-tracking complex.** (A) TipAct single-molecule dwell times on actin filaments. High temporal resolution kymograph of TipAct molecules showing multiple binding and unbinding events at a surface-bound actin filament (left), and a normalized distribution of TipAct dwell times (right), including single-exponential fit (red curve) which yielded an average dwell time of 111±6 ms (n=301) and corresponds to a single-molecule off-rate of about 0.01 $s^{-1}$. (B) FRAP experiments to probe protein off-rates at microtubule plus-ends. We analyze the recovery curves for mCherry-EB3 (top) and GFP-TipAct (bottom) at both free (red) and actin-bound (blue) microtubule tips. The solid lines show fits (see methods description below), and the thin colored lines and shaded areas show the average curves and SEM for n = 19 (free) and n = 13 (actin-bound) recovery profiles for mCherry-EB3, and n = 7 (free) and n = 6 (actin-bound) recovery profiles for GFP-TipAct, respectively. Note that the maximum recovery intensity of GFP-TipAct at actin-bound microtubule tips reaches values larger than one due to the variable amount of GFP-TipAct that can localize to the F-actin bundles independently of microtubules. (C) Schematics to interpret the FRAP experiments. Top, showing the unbinding rates for EB3 and TipAct at free microtubule plus ends. Bottom, showing the unbinding rates for EB3 and TipAct at actin-bound microtubule plus ends.

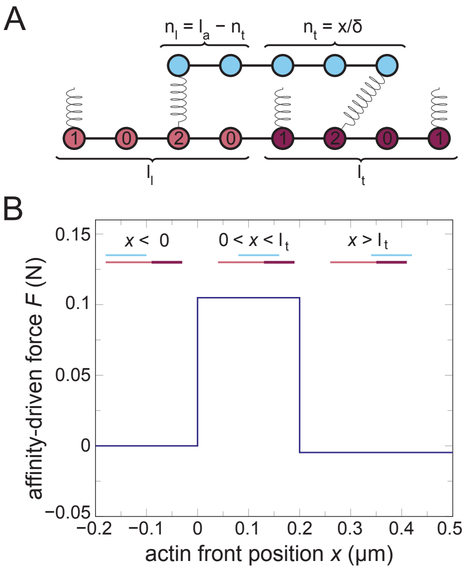


**Figure 8: Analytical calculations of the condensation forces.** (A) Parameter definitions: $\ell_\ell$ is the number of binding sites in the microtubule lattice region, and $\ell_t$ is the number of binding sites in the microtubule tip region. The actin filament contains $\ell_a$ binding sites, which are split into $n_\ell$ sites that overlap with the microtubule lattice and $n_t$ sites that overlap with the microtubule tip. The actin position $x$, defined as the distance between the front of the actin filament and the back of the microtubule tip region (see main Fig. 3C), equals $n_t \times \delta$, where $\delta$ is the lattice spacing between binding sites. We describe the full configuration of all cross-linkers by labeling each microtubule binding site with its binding state: no cross-linker is bound (0), a dangling cross-linker is bound (1), or a cross-linker is bound that connects with the actin (2). To calculate the condensation force, we take into account all possible extents to which the cross-linker can stretch. (B) The calculated condensation force on the actin filament as a function of the actin position $x$. When the actin filament is behind the tip ($x < 0$, top-left cartoon), the condensation force vanishes. When the actin filament is partially overlapping with the microtubule tip region ($0 < x < l_t$, top-center cartoon), there is a forward affinity driven force caused by the cross-linker affinity for the microtubule tip region compared to the lattice region. When the actin filament is sticking out in front of the microtubule ($x > l_t$, top-right cartoon), a tiny negative force acts on the actin filament due to the cross-linker affinity for the microtubule lattice region.

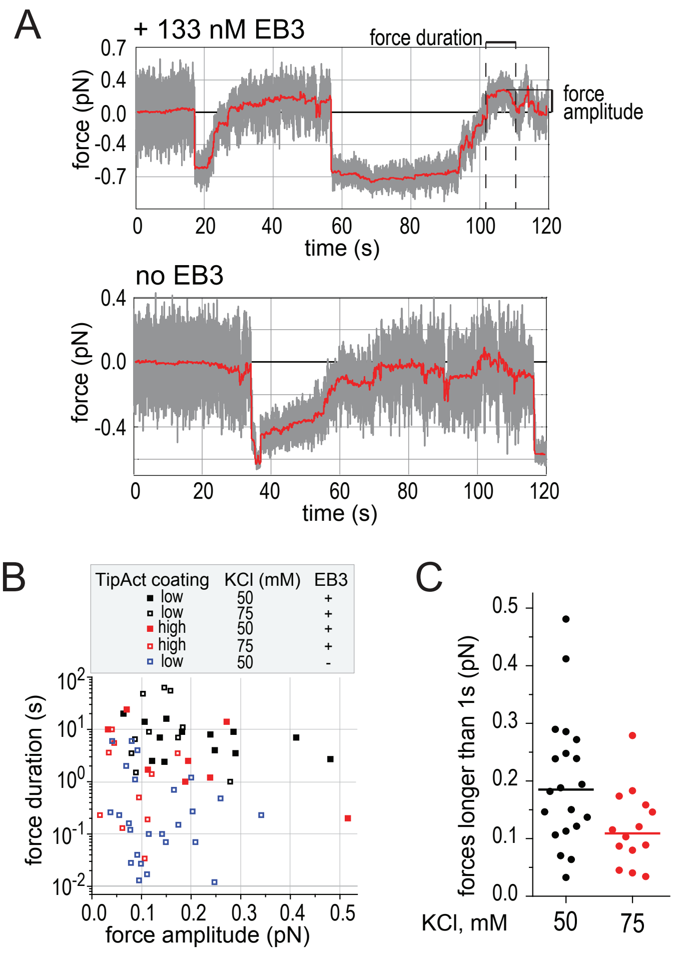


D

| | with EB3 | | without EB3 |
|---|---|---|---|
| | 50 mM KCl | 75 mM KCl | 50 mM KCl |
| force amplitude (pN) | 0.21 (±0.13) | 0.11 (±0.06) | 0.13 (±0.08) |
| force duration (s) | 7.6 (±6.7) | 11.5 (±19.3) | 0.94 (±1.8) |
| number of beads (# beads with force) | 32 (16) | 23 (12) | 25 (8) |
| number of force events (# force events >1s) | 21 (20) | 20 (14) | 25 (8) |

**Figure 9: Additional optical tweezer force measurements.** (A) Examples of full recordings of TipAct-coated beads interacting with growing microtubule ends in presence or absence of EB3. The initial displacement of the bead (resulting in a negative force) is also observed in the absence of EB3, due to the small (electrostatic) affinity of TipAct to the microtubule, which does not (sufficiently) discriminate between tip and lattice. Therefore the forward condensation force (but not the initial negative displacement) is dependent on EB3. Presented traces correspond to the truncated force traces shown in Fig.5C in the Main Text. (B) Distribution of duration and amplitude of forces obtained in conditions indicated in the legend. In absence of EB3, most forward force events last shorter than 1 s. (C) Distribution of force amplitudes for force events lasting longer than 1 s, comparing two different salt concentrations, with average forces of $(0.20 \pm 0.11)$ pN for 50 mM KCl and $(0.12 \pm 0.07)$ pN for 75 mM KCl (mean±SD). For the actin transport assays, we used 75 mM KCl. Straight lines are the median forces (0.19 pN for 50 mM KCl and 0.11 pN for 75 mM KCl). (D) Statistics of analyzed force curves and number of different beads for optical tweezer measurements. The amplitudes of the forces that lasted longer than 1s in the presence of EB3 are also indicated in C (i.e., 20 and 14 events for 50 and 75 mM KCl, respectively). Beads were selected for experiencing force events, which are events where the bead moves against an opposing force that is generated by a growing microtubule (as indicated in Fig.5B in the main text). For force amplitude and duration, means ± SD of all the measured force events are shown.

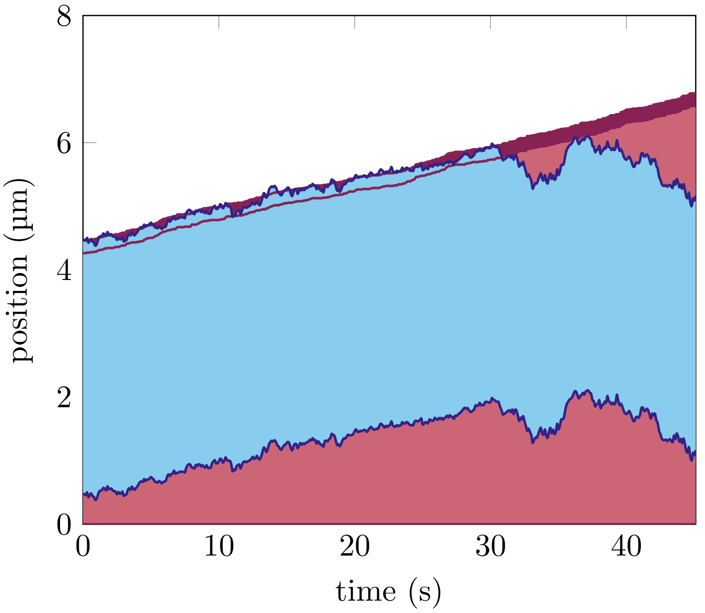


**Figure 10:** Simulations show that actin transport by growing microtubules also occurs when tip sites hydrolyse stochastically, resulting in an exponentially decaying density of tip sites. New sites are added with a rate of $6.25\,\mathrm{s}^{-1}$ (microtubule growth velocity of $v_g = 3\,\mu\mathrm{m}\,\mathrm{min}^{-1}$ from experimental data) and all tip sites decay to lattice sites with a rate of $0.25\,\mathrm{s}^{-1}$. These values lead to a density of tip sites that decays exponentially with a length scale of $0.2\,\mu\mathrm{m}$, matching the experimentally observed approximate tip length. The other parameter values are listed in Table 1.
